# Escape from Cell Uptake: Drug-Free Cancer Therapeutics Regulated Hydrophobicity and Negative Charge

**DOI:** 10.1101/2025.08.19.671160

**Authors:** Kazuki Moroishi, Masahiko Nakamoto, Satoshi Fujita, Kawahara Marie, Ryohei Katayama, Michiya Matsusaki

## Abstract

Self-aggregation and inducing cell membrane disruption in response to tumor microenvironment-stimuli is expected to be a promising approach for cancer treatment, but is limited by its insufficient stimuli-responsive cytotoxicity due to a lack of in-depth understanding of molecular characteristics, resulting in low selectivity of cell death induction. In this study, we focused on engineering polymer aggregation in detail to further improve tumor microenvironment-responsive cytotoxicity. PVA-U with grafting degrees (G.D.) of 3% (PVA-U3), 15% (PVA-U15), and 25% (PVA-U25) were synthesized and their aggregation properties cytotoxicity was evaluated. The difference in half maximal inhibitory concentration (IC50) values between pH 7.4 and pH 6.5 for PVA-U15 was 4.3-fold, which was greater than that of PVA-U25 at 2.8-fold, suggesting that tumor microenvironment-responsive cytotoxicity could be regulated by controlling G.D. of UDCA. Interestingly, PVA-U15 formed aggregates in the pericellular environment and adsorbed on the cell, effectively inducing cell death whereas PVA-U3 and PVA-U25 showed internalization in the cell. These results indicated that the balance of the surface charge and hydrophobicity could contribute to the adsorption on the cell membrane. These findings are expected to contribute to the development of membrane disruption strategies to control the aggregation properties and cell membrane interaction.

**Table of Contents:** Cancer therapeutics strategy targeting tumor microenvironment and cell adsorption of PVA-U with varying granting degree (G.D.). PVA-U is expected to aggregate and disrupt cell membrane in response to tumor microenvironment. PVA-U could regulate cell membrane adsorption by its negative charge and hydrophobicity due to controlling G.D. of UDCA, resulting in optimized tumor microenvironment selectivity.

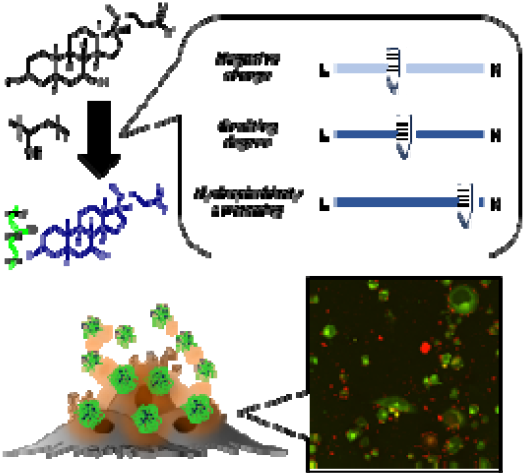

## 1. Introduction

Improvement of tumor selectivity is a key strategy in cancer therapy. For decades, drug delivery systems (DDS) have been widely studied to improve the tumoral accumulation of anticancer drugs.^1^ Sub-100 nm nanomaterials are mainly used in DDS, achieving selective penetration of drugs at the tumor site due to the enhanced permeability and retention (EPR) effect.^2^ However, the technique still has limited therapeutic efficiency because DDS strategies often target the nucleus to utilize anticancer drugs, requiring intracellular delivery through a multistep process.^3^ These processes cause insufficient induction of cell death. Furthermore, acquisition of drug resistance in cancer cells due to repeated chemotherapy is also one of the reasons for low therapeutic efficacy.^4^

In recent years, drug-free strategies aimed at overcoming these challenges have attracted attention. These strategies use nanomaterials that induce cell death through cell membrane disruption,^5,6^ mitochondrial dysfunction^7^, and reactive oxygen species (ROS) stress^8^ without anticancer drugs. They are expected to not only induce cell death with fewer processes, but also be effective against drug-resistant cells.

To further improve tumor selectivity, size-tuning of the nanomaterial is important. Although nanoscale aggregate is known to effectively penetrate and remain in the tumor, the size of the aggregate at the tumor site is controversial. Smaller sizes of nanoparticle with a diameter ~30 nm are better to circulate in the blood and penetrate tumor tissue than sub-100 nm nanoparticles.^9^ However, there is a contradictory problem in that larger nanoparticles are needed to enhance retention in tumor tissue. To address this issue, designs of nanomaterials that self-aggregate triggered by tumor microenvironment stimuli such as enzymes,^10,11^ pH,^12^ and light^13^ have recently been reported. This strategy will be expected to contribute not only to high penetration but also to further accumulation by reducing the backflow of aggregates from the tumor site into blood vessels.^14^

We have previously reported the development of a molecular block (MB) that self-aggregates and disrupts cancer cell membranes in the weak acid condition of a tumor microenvironment.^15–17^ The most recent MB consisted of a biocompatible poly(vinyl alcohol) (PVA) functionalized with ursodeoxycholic acid (UDCA), which is the most suitable bile acid with high weak acidic-sensitive cytotoxicity (Figure 1a).^17^ It was designed to be dispersed in the blood in nanoscale aggregates for efficient circulation and penetration into tumor tissue by the EPR effect. In the tumor microenvironment, where the pH is weakly acidic (pH 6.5-6.8),^18^ the MB induces selective cytotoxicity by aggregation due to an increase in hydrophobicity by protonation of UDCA. However, the difference of the half maximal inhibitory concentration (IC_50_) values between pH 7.4 and 6.5 in the MB was 2.1-fold, requiring further improvement in pH-sensitivity. This poor sensitivity by functionalization to polymer is possibly due to the burial of the UDCA moiety in the aggregates at pH 6.5, reducing the amount of UDCA that interacts with the cell membrane. Therefore, we need to focus on engineering aggregation and investigate its physical property in detail to further improve the tumor microenvironment-sensitivity. It has been reported that physicochemical characteristics such as size, surface charge, shape and hydrophobicity play important roles in interaction with the cell membrane and cellular internalization.^19^ Despite the many physicochemical arguments for improved tumor selectivity, many reports using DDS and self-aggregation strategies assume that most of these molecules would interact with cell membrane as they designed, and no reports have designed these properties in detail and regulated cell membrane interaction in these strategies.^10–13^

**Figure 1.**
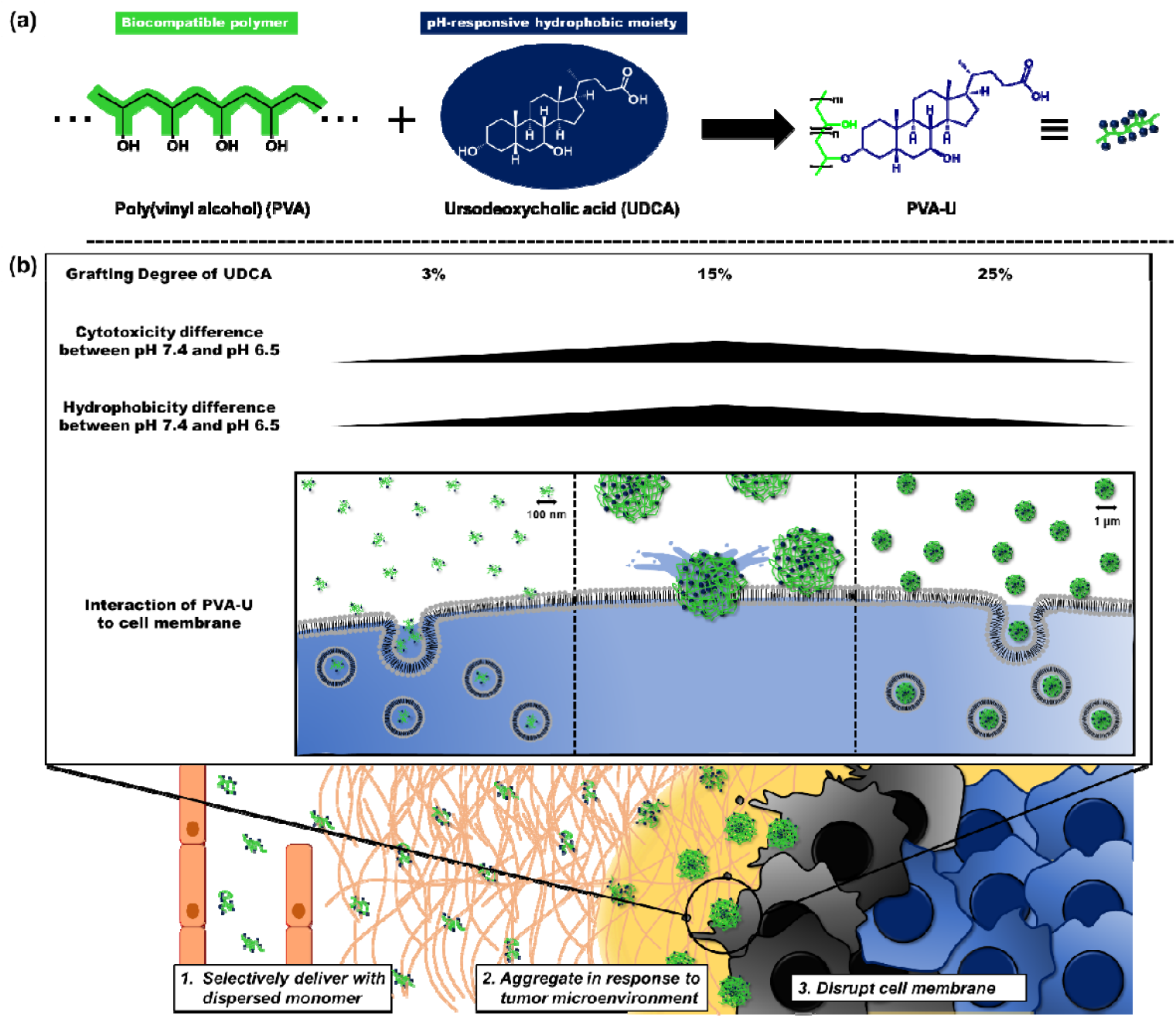
Overview of this study. (a) Chemical structure of PVA-U. A PVA-U was consisted of biocompatible poly(vinyl alcohol) and ursodeoxycholic acid (UDCA). (b) Schematic illustration of cancer therapeutics strategy targeting tumor microenvironment and cell adsorption of PVA-U with varying granting degree (G.D.). PVA-U was dispersed in blood with small size whereas this polymer was expected to aggregate in response to tumor microenvironment (pH 6.5) and subsequently induce cancer cell death by cell membrane disruption. PVA-U could regulate adsorption to cell membranes by its negative charge and hydrophobicity due to controlling G.D. of UDCA, resulting in optimized tumor microenvironment selectivity.

In this study, we designed the optimal aggregation behavior of PVA functionalized UDCA (PVA-U) by varying the grafting degree (G.D.) of UDCA for improvement of pH-responsive cytotoxicity. PVA-U with G.D. of 3% (PVA-U3), 15% (PVA-U15), and 25% (PVA-U25) were developed and their physical properties were evaluated. Although these polymers apparently showed pH-responsive aggregation, further investigation revealed that each polymer differed in its aggregation, surface charge, hydrophobicity, and morphology, depending on their G.D. The IC_50_ values of PVA-U15 at pH 7.4 were approximately 4.3-fold higher than that at pH 6.5, which is a much larger difference than that of PVA-U25; 2.8-fold. Moreover, most of the PVA-U15 was distributed in the pericellular environment and subsequently induced cell death whereas PVA-U3 and PVA-U25 internalized in the cell, suggesting these polymers could suppress cell death induction by cell uptake (Figure 1b). These results suggested that optimization of aggregation property and control of the interaction between the polymer and cell membrane are important factors in designing nanomaterials with highly pH-responsive cytotoxicity.

## 2. Result and discussion

### 2.1. Synthesis of PVA-U

A Poly (vinyl alcohol) functionalized by ursodeoxycholic acid (PVA-U) was considered using PVA as a biocompatible polymer and UDCA as a trigger moiety of aggregation and cell membrane disruption in response to a weak acidic condition (Figure 1a). UDCA was conjugated to PVA via mesylation according to our previously reported procedure (Scheme S1).^17^ Mesylation of UDCA was confirmed by the shift in the peak assigned **e** from 3.6 ppm to 4.6 ppm and emergence of a proton from methansulfonyl group in proton nuclear magnetic resonance (^1^H NMR) (Figure S1). To synthesize PVA-U with different G.D., UDCA was reacted in 2.5, 5, and 10 equivalents per unit mole of PVA. ^1^H NMR characterization confirmed the G.D. of UDCA in each polymer was 3%, 15%, and 25% (Figure S2-S4). Each concentration of UDCA against 1 mg mL^−1^ PVA-U was estimated at 0.481, 1.44, and 1.76 mM, respectively (Table S1). Hereafter, the PVA-U with G.D. of 3%, 15%, and 25% are named PVA-U3, PVA-U15, and PVA-U25, respectively. PVA (PVA-U0) was also used in their characterization as a control.

### 2.2. pH-responsiveness of PVA-U

A PVA-U is expected to aggregate in response to weak acidic pH due to an increase in hydrophobicity by protonation of UDCA. First, to confirm weak acidic pH-responsiveness, p*K*_a_ values of PVA-U3, PVA-U15, and PVA-U25 were evaluated by pH titration. The titration curve of PVA-U also showed a buffer effect at between pH 8.5 and 5.0, confirming weak acidic pH-responsiveness of PVA-Us (Figure S5a). The p*K*_a_ values of PVA-U3, PVA-U15, and PVA-U25 were 6.34, 6.86, and 6.40, respectively, which were all higher than that of UDCA (Figures 2a and S5b). This is because a hydrophobic moiety would be formed in the PVA-U aggregates as the pH decreased, possibly reducing the exchange of protons of the carboxyl group. Therefore, the buffer effect of PVA-Us would appear to terminate at a higher pH as the p*K*_a_ of UDCA increased in a concentration-dependent manner (Figure S5b). These polymers showed higher p*K*_a_ values than UDCA likely due to suppression of proton exchange of UDCA within aggregates as the pH decreased, as also seen in the case of UDCA.

**Figure 2.**
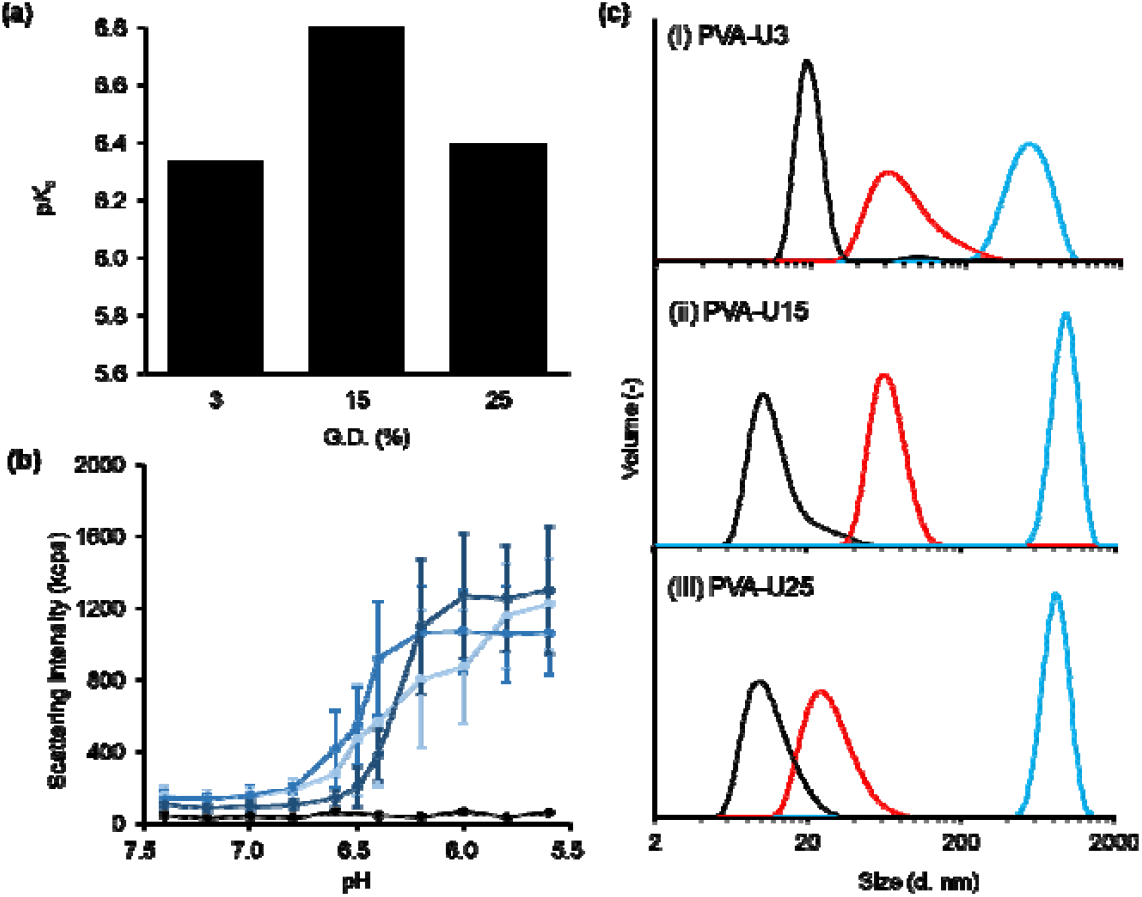
pH-responsive aggregation property of PVA-Us with different grafting degree. (a) p*K*_a_ value of each PVA-U. (b) pH-dependent scattering intensity of PVA-U0 (black), PVA-U3 (light blue), PVA-U15 (blue), and PVA-U25 (dark blue) in PBS at a concentration of 10 µg mL^−1^ immediately after preparation at 37 °C. (c) Size distribution of PVA-U3 (i), PVA-U15 (ii), and PVA-U25 (iii) at pH 7.4 (black), pH 6.5 (red), and pH 5.6 (blue) immediately after preparation at 37 °C.

Next, the aggregation behavior of PVA-Us in response to a weak acidic condition was evaluated using scattering intensity by dynamic light scattering (DLS) immediately after preparation. PVA-U0 showed less than 100 kilos count per second (kcps) of scattering intensity at each pH, suggesting no pH-responsive aggregation (Figure 2b). In contrast, although the scattering intensity of PVA-Us was less than 200 kcps at pH 7.4, these values began to increase from pH 6.8, and was over 1,000 kcps at below pH 6.0. These results revealed that PVA-U aggregated in response to a weak acidic condition due to an increase in its hydrophobicity by protonation of UDCA. To analyze the pH response of a material, the slope of the fitting curve for a given parameter against pH should be determined.^24^ Based on this, we performed further analysis of the aggregation behavior of PVA-U3, PVA-U15, and PVA-U25 using the graph of the scattering intensity of each polymer against pH (Figure S6a-c). Two parameters were used in this analysis; (1) aggregation pH: the pH at which the scattering intensity begins to increase and (2) the slope from an increase in scattering intensity until it reaches a plateau. The aggregation pH of PVA-U3 and PVA-U15 were pH 6.7 whereas that of PVA-U25 was pH 6.6 (Figure S6d). PVA-U25 started to aggregate at a slightly lower pH than PVA-U3 and PVA-U15 although these differences were not significant. Regarding the slope, it was slightly lower in PVA-U3 than in PVA-U15 and PVA-U25 (Figure S6e). However, these differences were not significant, which each PVA-U showed a similar aggregation property. In addition, the aggregation ratio, which is the ratio of scattering intensity normalized by each at pH 5.6, against the ionization ratio of PVA-U by titration was calculated. Each aggregation ratio started to increase at around 50 ~ 60% ionization, revealing that each PVA-U initiated aggregation at each p*K*_a_ value (Figure S7). The size distribution of PVA-Us also indicated pH-responsive aggregation; approximately 20, 60, and 1,000 nm at pH 7.4, 6.5, and 5.6, respectively (Figure 2c). These results suggested that PVA-Us started to aggregate around their p*K*_a_, suggesting that the aggregation is due to protonation of UDCA.

### 2.3. Aggregation and surface physical property of PVA-Us

Surface physical property is important to understand the aggregation property and cytotoxicity of PVA-Us. The surface charge of PVA-Us was evaluated by measurement of zeta potential. The zeta potentials of PVA-U0, PVA-U3, PVA-U15, and PVA-U25 at pH 7.4 were −1.92, −16.3, −22.5, and −26.2, respectively, with values decreasing with an increasing G.D. due to an increase in the unit amount of ionized UDCA (Figure 3a). In contrast, the values at pH 6.5 were 1.62, −14.4, −20.5, and −23.6, respectively. Each zeta potential at pH 6.5 was higher than that at pH 7.4 and decreased with an increase in G.D. as at pH 7.4. Moreover, the zeta potential of these polymers at pH 5.6 showed higher values than those at pH 6.5 (Figure S8). These results revealed that the surface charge of PVA-U aggregates would increase due to further protonation of UDCA. Hydrophobicity of PVA-Us was investigated by contact angle (CA) measurement of 1x PBS droplet on surfaces coated with 10 mg mL-1 PVA-Us solution in DMSO. The CA test is generally used for materials where the hydrophobicity changes in response to stimuli.^25^ The PBS droplet at pH 6.5 on the PVA-U surface showed higher water repellency than that at pH 7.4 (Figures 3b and S9). The CA of a PBS droplet at pH 6.5 on the PVA-U3, PVA-U15, and PVA-U25 were 40.7 °, 61.5 °, and 72.5 °, respectively, which were higher than those at pH 7.4: 37.1 °, 46.9 °, and 68.1 °, respectively, due to an increase in hydrophobicity by protonation of UDCA (Figure 3c). CA values also increased with an increase in G.D. due to an increase in the amount of UDCA units in a polymer. Notably, the CA difference between pH 7.4 and pH 6.5 of PVA-U3, PVA-U15, and PVA-U25 were 3.60, 14.6, and 4.37, with PVA-U15 showing the largest difference in hydrophobicity (Figure S10). This result suggested that PVA-U15 would have high hydrophilic-hydrophobic switching capacity. This finding was reasonable according to the difference in hydrophobicity change of the temperature-responsive materials previously reported.^26^

**Figure 3.**
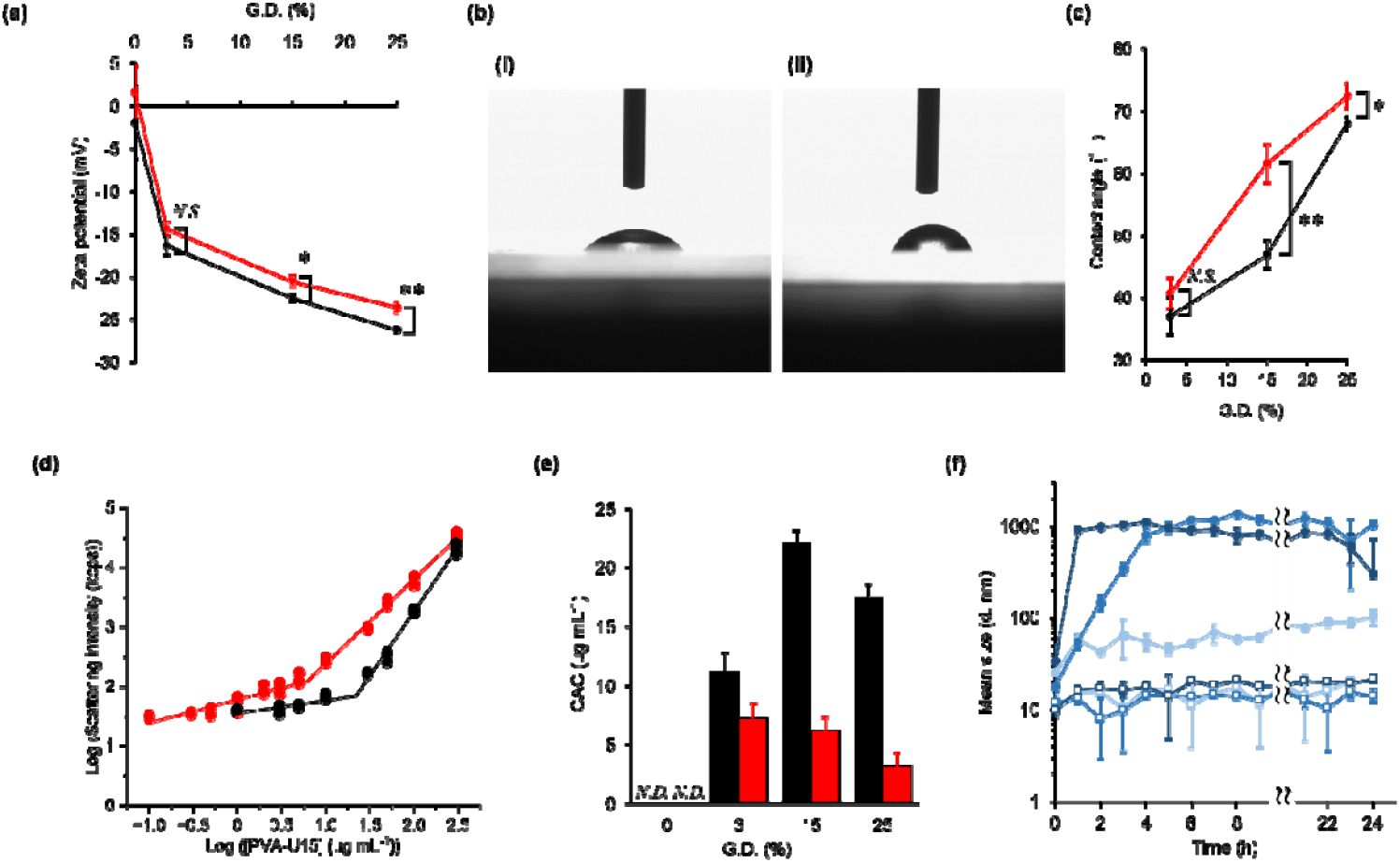
Surface physical property and concentration and time dependent aggregation of PVA-Us. (a) Zeta potential values of PVA-Us at pH 7.4 (black) and pH 6.5 (red) immediately after preparation at 37 °C (n = 3). (b) Images of droplet of 1X PBS at pH 7.4 (i) and pH 6.5 (ii) to slide grass coated by PVA-U15. (c) Contact angle of 1X PBS on the surface of PVA-U3, PVA-U15, and PVA-U25 at pH 7.4 (black) and pH 6.5 (red) (n = 3). (d) Concentration-dependent light scattering intensity of PVA-U15 in PBS at pH 7.4 (black) and pH 6.5 (red) immediately after preparation at 37 °C. (e) CAC values of PVA-Us at pH 7.4 (black) and pH 6.5 (red) (n = 3). (f) Time-dependent mean size of PVA-U3 (light blue), PVA-U15 (blue), and PVA-U25 (dark blue) at pH 7.4 (square) and pH 6.5 (circle) incubated for 24 hours at 37 °C (n = 3). In (e), error bars represented fitting error. Statistical analysis was performed using unpaired two-tailed Student’s *t*-test. Data are presented as mean ± S.D. (*N*.*S*.: no significant difference, ^*^*p* < 0.05, ^**^*p* < 0.01, *N*.*D*.: not determined).

To further understand pH-responsive aggregation property, we evaluated the critical aggregation concentration (CAC) of PVA-U by measuring scattering intensity using DLS.^17,22^ When the solution reaches the CAC, the scattering intensity of DLS increases suddenly due to the formation of aggregates.^22^ We determined the CAC using a piecewise fitting model which can calculate the critical point. The scattering intensity of PVA-U0 at pH 7.4 and pH 6.5 were comparable and both values showed a slight concentration-dependent increase, at most 100 kcps (Figure S11a). The sizes of PVA-U0 at a concentration of 300 µg mL^−1^ at pH 7.4 and pH 6.5 were 12 nm and 15 nm, respectively, which are comparable values to 5k poly(ethylene) glycol that is a dispersion (Figure S12). This result indicated that although the critical point was observed at both pHs (Figure S11a), PVA-U0 was considered dispersed at concentrations below 300 µg mL^−1^. In contrast, the size of each PVA-U at a concentration of 300 µg mL^−1^ was more than 20 nm, which is significantly higher than 5k poly(ethylene) glycol. Moreover, each scattering intensity of PVA-U showed a marked increase in scattering intensity with increasing concentration and exceeded 1,000 kcps at 300 µg mL^−1^ (Figures 3d, S11b, and S11c), suggesting aggregate formation. CAC values of PVA-U3, PVA-U15, and PVA-U25 were 11.2, 22.1, and 17.4 µg mL^−1^ at pH 7.4, and 7.20, 6.20, and 3.14 µg mL^−1^ at pH 6.5, respectively (Figure 3e). These results suggested that PVA-Us at pH 6.5 self-aggregated at a lower concentration than those at pH 7.4, revealing that their aggregation properties significantly increased in response to a weak acidic condition by protonation of UDCA. In comparison to CAC values at pH 7.4, that of PVA-U3 was the lowest of the PVA-Us. This is because the zeta potential of PVA-U3 was higher than that of PVA-U15 and PVA-U25 due to a lesser amount of ionized UDCA (Figure 3a), suggesting that the polymers interact easily with each other and form small aggregates even at low concentration. In contrast, PVA-U15 and PVA-U25 showed less aggregation below CAC due to electron repulsion between polymers by higher anion charge than PVA-U3. CAC values of PVA-Us at pH 6.5 decreased with an increase of G.D. due to an increase in hydrophobicity (Figure 3c), demonstrating that aggregation occurs easily at low concentration. Thus, the pH-responsive aggregation property could be tuned by the balance between its charge and hydrophobicity based on G.D. The size of PVA-Us was measured every 1 h at a concentration of 10 mg mL^−1^, which is a CAC between pH 7.4 and pH 6.5. The size of PVA-U3, PVA-U15, and PVA-U25 at pH 7.4 stayed at approximately 20 nm until 24 h incubation (Figures 3f and S13) while the PVA-U3 at pH 6.5 gradually increased over time, forming approximately 100 nm aggregates after 24 h. PVA-U15 and PVA-U25 reached a plateau in size after 4 and 1 h, respectively, forming aggregates of around 1,000 nm. These results revealed that the aggregation rate was faster as the hydrophobicity increased due to an increase in G.D.

Taken together, the pH-responsive aggregation property of PVA-U was more sensitive with an increase in G.D. at pH 6.5. However, the hydrophilic-hydrophobic switching capacity of the aggregates was found to be maximized at an intermediate G.D.: PVA-U15, suggesting the usefulness of tuning their G.D.

### 2.4. Morphology of PVA-U

To understand the morphology of each PVA-U aggregate, we observed PVA-U3, PVA-U15, and PVA-U25 aggregates at pH 7.4 and pH 6.5 by transmission electron microscopy (TEM). Each of the PVA-U aggregates at pH 6.5 were larger than those at pH 7.4, confirming pH-responsive aggregation (Figures 4a-4f). The sizes of PVA-U3, PVA-U15, and PVA-U25 at pH 7.4 were less than 200 nm, 100 nm, and 300 nm, respectively, suggesting the smallest size for PVA-U15 (Figures 4g and S14). In contrast, those at pH 6.5 were less than 600 nm, 1,100 nm, and 500 nm, respectively. These results revealed that PVA-U15 showed the largest size difference between pH 7.4 and pH 6.5. Although the size distribution of PVA-U3 and PVA-U15 from TEM images were a good agreement with each of the DLS results, that of PVA-U25 at pH 6.5 showed a smaller size distribution than that of the DLS result. It is possible that the shrinkage of the aggregates in the TEM images was due to the freeze-drying, compared to the DLS results which were measured in solution. PVA-U3 formed relatively spherical and well-ordered aggregates at pH 6.5 whereas PVA-U25 showed disordered aggregates, suggesting PVA-U aggregates would be disordered with an increase in G.D. (Figures 4d-4f). To confirm these tendencies, the circularity index of each PVA-U was quantified from TEM images (Figure 4h). The indices of PVA-U3, PVA-U15, and PVA-U25 at pH 7.4 were approximately 0.73, which showed comparable circularity. In contrast, those at pH 6.5 were 0.73, 0.56, and 0.42, with circularity decreasing with an increase in G.D. Notably, PVA-U15 and PVA-U25 showed significant differences in circularity between pH 7.4 and pH 6.5. This suggests that PVA-U formed disordered aggregation due to fast aggregation as mentioned in the DLS result (Figure 3f). Therefore, PVA-U25 aggregated rapidly within 2 h, forming disordered aggregation whereas PVA-U3 formed ordered aggregates by slow aggregation.

**Figure 4.**
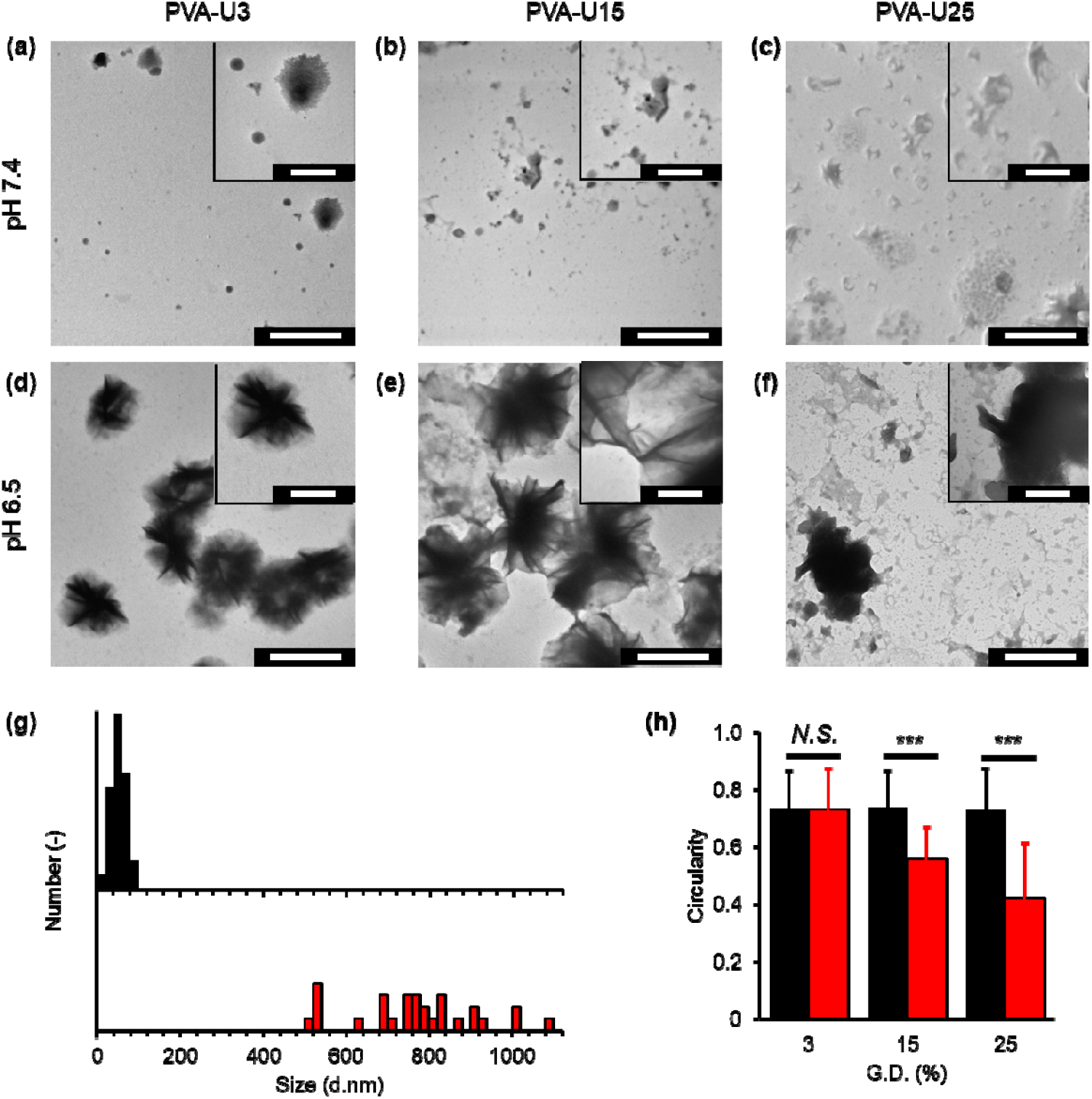
TEM images of PVA-U3, PVA-U15, and PVA-U25 aggregates incubated for 6 h in 1X PBS at pH 7.4 (a-c) and pH 6.5 (d-f). Scale bars for images and enlarged images are, respectively, 1000 nm and 500 nm. (g) Size distribution of PVA-U15 aggregates at pH 7.4 (black) and pH 6.5 (red) analyzed from TEM images. (h) Circularity of PVA-U3, PVA-U15, and PVA-U25 aggregates at pH 7.4 (black) and pH 6.5 (red) analyzed from TEM images. In (g) and (h), 30 aggregates in the images were randomly selected for analysis. Statistical analysis was performed using unpaired two-tailed Student’s *t*-test. Data are presented as mean ± S.D. (*N*.*S*.: no significant difference, ^***^*p* < 0.001).

To summarize the physical properties of each PVA-U, these polymers dispersed in solution at a pH above each p*K*_a_ (6.3 ~ 6.9) by ionized UDCA whereas they formed aggregates at a pH below their p*K*_a_ due to hydrophilic-hydrophobic switching by protonation of UDCA. Further evaluation of the physical properties also revealed that the surface properties of aggregates such as hydrophobicity and zeta potential, morphology, and aggregation rate depended on their G.D. Specifically, PVA-U3 showed less hydrophobicity and negative charge than the other PVA-Us at both pHs, resulting in low pH-responsive aggregation property and slowly formed ordered aggregates whose sizes were approximately 100 nm at pH 6.5 after 24 h (Figure 5). In PVA-U15, although the surface charge of aggregates at pH 6.5 was more positive than those at pH 7.4, the charge at both pHs was more negative than that of PVA-U3. Furthermore, these polymers aggregated to 1,000 nm in size after 4 h at pH 6.5, quickly forming larger aggregates than PVA-U3. Notably, the hydrophobicity of PVA-U15 at pH 6.5 significantly increased compared to that at pH 7.4, suggesting higher hydrophilic-hydrophobic switching capacity than the other PVA-Us. Given amphiphilicity is the key factor for cell membrane disruption, its high hydrophilic-hydrophobic switching capacity would be a great contribution to pH-responsive cytotoxicity. PVA-U25 was the most negatively charged and hydrophobic of the PVA-Us at both pHs, and formed disordered aggregates of 1,000 nm after 1 h at pH 6.5. Hence, the surface charge, hydrophilic-hydrophobic switching capacity, aggregation size, rate, and order of PVA-U could be tailored by controlling the G.D. of UDCA.

**Figure 5.**
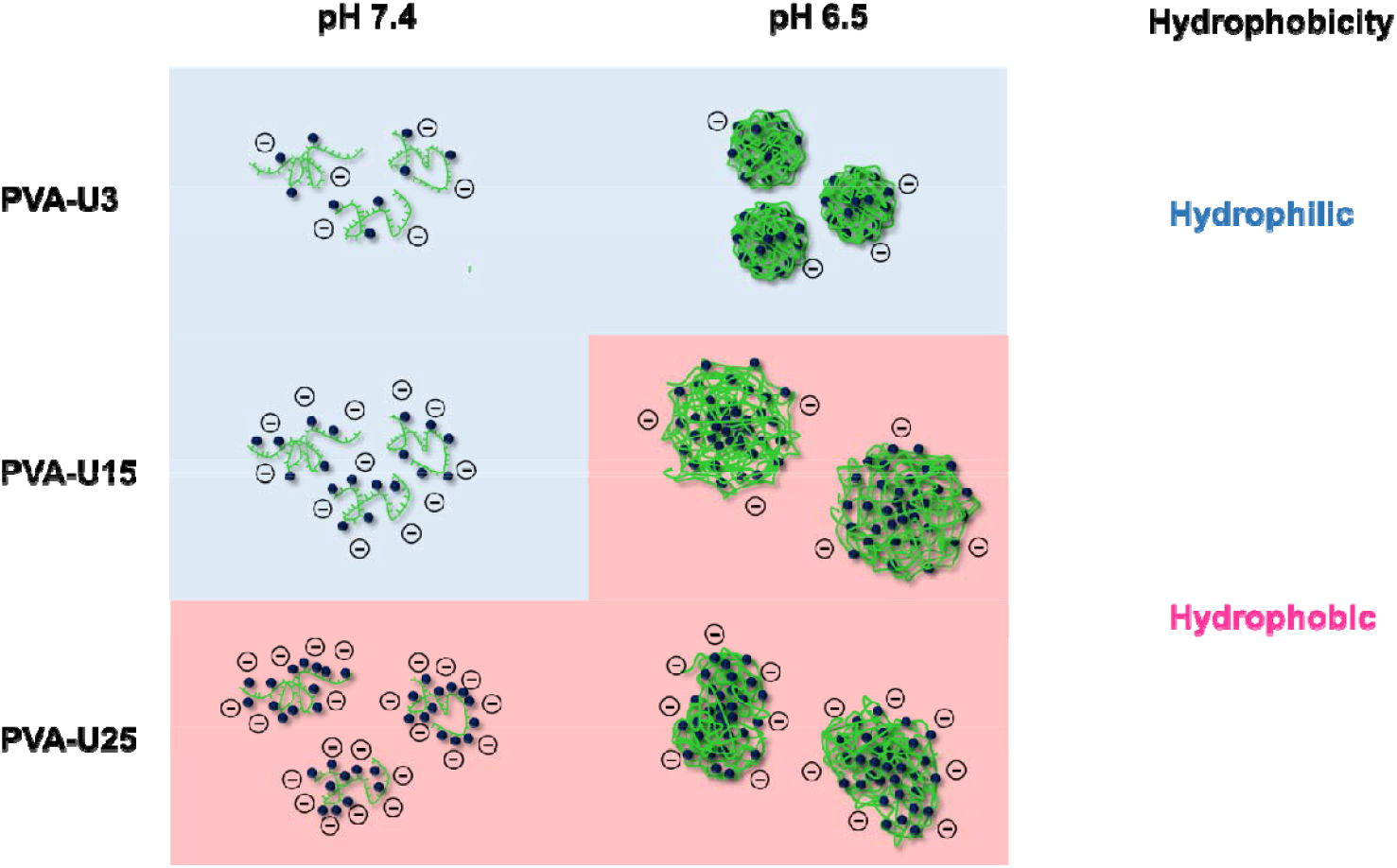
Schematic illustration of physical property of PVA-U. Each PVA-U aggregated in response to weak acidic condition. PVA-U3 was more hydrophilic, higher zeta potential at both pHs, and more ordered shape at pH 6.5 compared to other PVA-Us due to slow aggregation formation. PVA-U15 forms aggregates of approximately 1 µm in 4 hours and is more disordered than PVA-U3 at pH 6.5. Notably, PVA-U15 showed markedly increase in hydrophobicity in response to pH, suggesting high hydrophilic-hydrophobic switching capacity. PVA-U25 was more hydrophobic, lower zeta potential at both pHs, and less ordered shape at pH 6.5 compared to other PVA-Us due to fast aggregation formation. As a result, PVA-U showed higher hydrophobicity, lower zeta potential, and less ordered shape of aggregates with an increase in grafting degree.

### 2.5. pH-responsive cytotoxicity of PVA-Us

Tumor microenvironment responsive cytotoxicity of PVA-Us against pancreatic cancer (MIAPaCa-2) cells was evaluated. 50% inhibition concentration (IC_50_) values of PVA-U0, PVA-U3, PVA-U15, and PVA-U25 were calculated by measuring the cell viability of MIAPaCa-2 cells treated with each polymer after incubation for 24 h (Figure 6a). The IC_50_ values of UDCA at pH 7.4 and 6.5 have previously reported to be approximately 1,800 µm and 240 µm, respectively.^17^ The shape of MIAPaCa-2 cells treated with PVA-U0 did not change at a concentration of 50 µg mL^−1^ (Figure S15) and the cell viability of MIAPaCa-2 cells treated with PVA-U0 at pH 7.4 and pH 6.5 were more than 80% (Figures S16a and S16b). The shape of MIAPaCa-2 cells treated with PVA-U3 also did not change and the cell viability at pH 7.4 and pH 6.5 exceeded 90% and 70%, respectively (Figures S16c and S16d). The IC_50_ values of these polymers could not be calculated due to their high biocompatibility, which did not induce tumor microenvironment responsive cytotoxicity. In contrast, MIAPaCa-2 cells treated with PVA-U15 and PVA-U25 showed cell elongation at pH 7.4 whereas cells shrunk at pH 6.5 (Figure S14) and their cell viability decreased with an increase in concentration (Figures S16e-S16h). IC_50_ values of PVA-U15 and PVA-U25 at pH 7.4 were 1.3 µM and 1.6 µM whereas those at pH 6.5 were 0.31 µM and 0.55 µM, respectively, suggesting tumor microenvironment responsive cytotoxicity by cell membrane disruption. These results are due to an increase in hydrophobicity by protonation of UDCA. In addition, the IC_50_ values at pH 7.4 based on UDCA concentration were 180 µM and 380 µM at pH 7.4, and 43.0 µM and 130 µM at pH 6.5, respectively (Figure S17). IC_50_ values of PVA-U15 and PVA-U25 at both pHs were lower than that of UDCA, suggesting an increase in local concentration of UDCA by conjugating to PVA. Differences in the IC_50_ values of PVA-U15 and PVA-U25 between pH 7.4 and 6.5 were 4.3-fold and 2.8-fold, revealing higher pH-responsive cytotoxicity of PVA-U15. Although this value was lower than UDCA molecules: 7.5-fold, PVA-U15 induced cell death at a lower concentration than UDCA molecules and would be expected to selectively deliver to tumor tissue by EPR effect *in vivo* due to conjugation to PVA.

**Figure 6.**
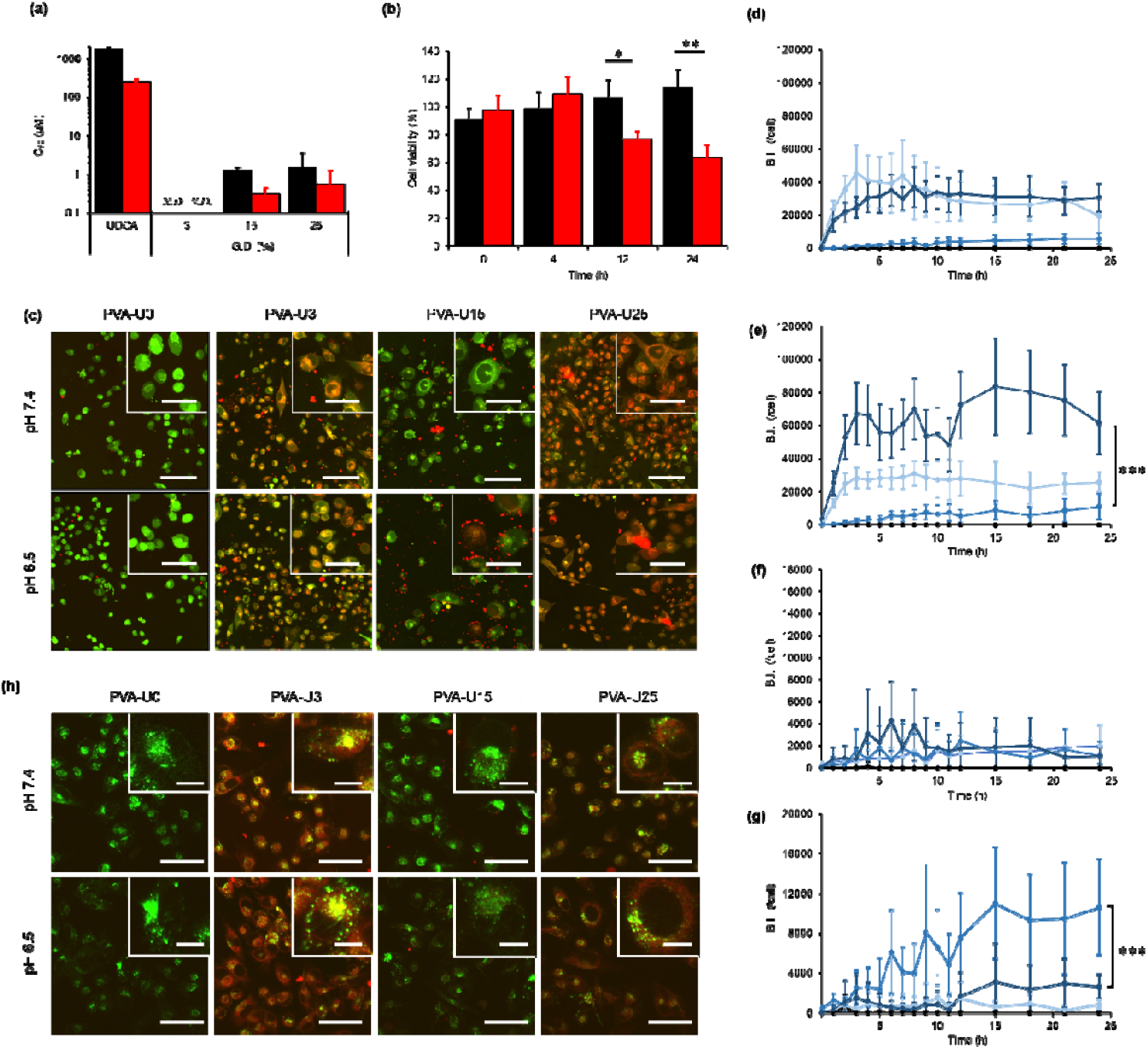
Cytotoxicity and Cell adsorption property of PVA-U varying G.D. (a) IC_50_ values of MIAPaCa-2 cells treated with UDCA, PVA-U3, PVA-U15, and PVA-U25 for 24 h incubation at pH 7.4 and 6.5 (n = 3). (b) Cell viability of MIAPaCa-2 cells treated with 10 µg mL^−1^ PVA-U15 at pH 7.4 and pH 6.5 for 0, 4, 12, and 24 h incubation (n = 3). (c) Confocal images of MIAPaCa-2 cells treated with Cell tracker deep red and 10 µg mL^−1^ PVA-U0-R, PVA-U3-R, PVA-U15-R, and PVA-U25-R at pH 7.4 and pH 6.5 for 12 h incubation. Cell tracker deep red and PVA-Us were shown in green and red color, respectively. Scale bars of images and enlarge images are 100 µm and 50 µm, respectively. Quantification of intracellular fluorescence intensity of MIAPaCa-2 cells treated with PVA-U0 (black), PVA-U3 (light blue), PVA-U15 (blue), and PVA-U25 (dark blue) incubated for 24 h at pH 7.4 (d) and pH 6.5 (e) (n = 11). Quantification of pericellular fluorescence intensity of MIAPaCa-2 cells treated with PVA-U0 (black), PVA-U3 (light blue), PVA-U15 (blue), and PVA-U25 (dark blue) incubated for 4 h at pH 7.4 (f) and pH 6.5 (g) (n = 11). (h) Confocal images of MIAPaCa-2 cells treated with lysotracker and 10 µg mL^−1^ PVA-U0-R, PVA-U3-R, PVA-U15-R, and PVA-U25-R at pH 6.5 for 4 h incubation. Lysotracker deep red and PVA-Us were shown in green and red color, respectively. Scale bars of images and enlarge images are 50 µm and 10 µm, respectively. Statistical analysis was performed using unpaired two-tailed Student’s *t*-test. Data are presented as mean ± S.D. (*N*.*S*.: no significant difference, ^*^*p* < 0.05, ^**^*p* < 0.01, ^***^*p* < 0.001).

To confirm whether PVA-U induce cell death by disrupting cell membrane, we also measured the leakage of lactate dehydrogenase (LDH), an intracellular substance, in MIAPaCa-2 cells treated with each PVA-U. LDH leakage was little confirmed in PVA-U0, PVA-U3, and PVA-U15 at pH 7.4, while PVA-U25 showed a leakage of about 8% (Figure S18). PVA-U0 also showed little leakage of LDH at pH 6.5, as did pH 7.4. In contrast, PVA-U3, PVA-U15, and PVA-U25 showed significant higher leakage at pH 6.5 than that at pH 7.4. These result suggested PVA-U induce cell death by disrupting cell membrane. Notably, PVA-U15 showed the highest LDH leakage rate of approximately 30%, and the difference in LDH leakage rates between pH 7.4 and pH 6.5 was the largest among PVA-U, which was consistent results with the cytotoxicity experiments. Although the cell viability at the same concentration of PVA-U15 at pH 6.5 was 11%, the LDH leakage rate was not relatively high value, which may be due to the time needed for LDH to leak out into the medium after cell death is induced.

Cell viability of MIAPaCa-2 cells treated with PVA-U15, which showed the highest pH-responsive cytotoxicity, was measured over time. Cell viability at pH 7.4 was more than 100% after 24 h of incubation whereas these values at pH 6.5 started to decrease from 12 h and showed 63% after 24 h of incubation at a concentration of 10 µg mL^−1^ (Figure 6b). This value was in good agreement with the cell viability result in concentration-dependent cytotoxicity evaluation (Figure S16f).

To confirm how PVA-U with varying G.D. contributed to tumor microenvironment responsive cell death induction, we conjugated rhodamine B isothiocyanate (RBITC) to PVA-U and observed cell adsorption behavior of labeled PVA-U (PVA-U-R) to MIAPaCa-2 cells which stained with CellTracker™ deep red by confocal microscopy. PVA-U0-R did not show aggregation and cell adsorption at pH 7.4 and pH 6.5 (Figures S19 and S20) whereas PVA-U3-R showed aggregation at both pHs and co-localization of polymer and cytosol was observed (Figures 6c, S21, and S22). Although PVA-U15-R aggregated with each other at both pHs, these aggregates selectively gathered in the pericellular environment after 12 h of incubation and subsequently changed cancer cell shape at pH 6.5 compared to that at pH 7.4 (Figures S23 and S24). This result was in good agreement with a decrease in cell viability of MIAPaCa-2 cells at pH 6.5 from 12 h of incubation (Figure 6b). The cytosol color of MIAPaCa-2 cells treated with PVA-U25-R began to change from green to red at both pHs, suggesting PVA-U25 was more internalized into the cell than PVA-U3 (Figures S25 and S26). Interestingly, PVA-U25-R aggregates were not seen in the presence of cells despite of forming 1,000 nm aggregates within 1 h at pH 6.5 by DLS measurement (Figure 3f). We also quantified the intracellular and pericellular brightness intensity (B.I.) of each PVA-U. Intracellular B.I of PVA-U0-R showed little B.I. at either pH, which did not absorb on the cell (Figures 6d and 6e). In contrast, that of PVA-U3-R increased for 3 h and maintained a comparable B.I. at both pHs until 24 h of incubation. That of PVA-U25-R also increased to over 30,000 after 6 h at pH 7.4 and to 60,000 after 3h of incubation pH 6.5, respectively, maintaining its B.I. until 24 h of incubation. This result suggested PVA-U25 internalized to cancer cells in response to a weak acid condition. In contrast, the intercellular B.I. of PVA-U15-R was over 5,000 at pH 7.4 and over 10,000 at pH 6.5, respectively, which is significantly lower than that of PVA-U3-R and PVA-U25-R at each pH. In pericellular B.I., PVA-U0-R, PVA-U3-R, PVA-U15-R, and PVA-U25-R showed comparable B.I. at approximately 2,000 at pH 7.4, which was significantly lower than intracellular (Figure 6f). These values of PVA-U0-R, PVA-U3-R, and PVA-U25-R at pH 6.5 also showed less than 2,000, suggesting these polymers were less distributed in the pericellular environment than intracellularly (Figure 6g). In contrast, PVA-U15-R at pH 6.5 showed more than 10,000, which was much higher than that of other polymers. Interestingly, this result showed an opposite trend to the result of intracellular B.I., which means most of the PVA-U15 was distributed in the pericellular environment, resulting in the highest tumor microenvironment responsive cytotoxicity.

From the results of confocal observation, we hypothesized that cell uptake prevents PVA-U from efficiently inducing cell death. To confirm this hypothesis, we evaluated the cell uptake of PVA-Us by incubating MIAPaCa-2 cells treated with each PVA-U-R and lysotracker. PVA-U0-R did not show co-localization with lysotracker at either pH because PVA-U0-R did not aggregate as well as figure 6c (Figures 6h, S27, and S28). In PVA-U3, and PVA-U25, co-localization between PVA-U and lysotracker was observed at both pHs. The brightness intensity of PVA-U3 and PVA-U25 at each location was also in good agreement with that of each lysotracker in line scan analysis, suggesting these polymers were taken up in acidic vesicle in the cell (Figure S29). PVA-U3-R and PVA-U25-R were distributed not only in the lysosome but also the cytosol which was considered to be due to endosome escape by endosome membrane disruption during the decrease in pH in acidic vesicle lumen. In contrast, PVA-U15 did not show co-localization with lysotracker at either pH in confocal images and line scan analysis, suggesting cell uptake of PVA-U15 was not observed. Furthermore, we tried to inhibit the energy-dependent cell uptake by incubating MIAPaCa-2 cells treated with PVA-U-Rs under 4 °C. PVA-U-R did not internalize in the cell, which also suggested that the cell uptake of PVA-U3 and PVA-U25 was by energy-dependent cell uptake like the endocytosis pathway (Figure S30).

Based on these results, we discuss the cell interaction property of PVA-Us as follows. PVA-U0 did not adsorb on the cell membrane because PVA-U0 did not have UDCA which is a cell interaction moiety. PVA-U3 did not show cytotoxicity at either pH 7.4 or pH 6.5, although cellular uptake was observed. This may be due to the low negative charge and hydrophobicity of PVA-U3, as well as the cellular insertion ability of slightly modified UDCA, resulting in cell membrane adsorption (Figure 7a), Size-dependent cellular uptake is also possible at pH 6.5, since PVA-U3 formed aggregates of approximately 100 nm in size at pH 6.5 by DLS measurement (Figure 3f). These size ranges have been reported to be readily taken up by cancer cells (Figure 7b).^27^ However, these cell uptake remains unclear and requires further investigation. PVA-U15 at pH 7.4 had a higher anion charge than PVA-U3, exerting lower cell membrane interaction by electronic repulsion (Figure 7c) whereas that at pH 6.5 formed micro-scale aggregates in the pericellular environment without internalization in the cell, effectively adsorbing on the cell and subsequently inducing cell death by cell membrane disruption (Figure 7d). This result suggested to be due to its great hydrophobicity switching. Regarding the mechanism of inducing cancer cell death by self-aggregation and disruption of the cell membrane, it has been reported that induction cell death maybe due to forming transit pores by membrane perturbation^28^ due to an increase in local concentration of UDCA, and inducing cell rupture by targeting lipid rafts^29^. Therefore, PVA-U15 would be likely to exert cell membrane disruption by membrane perturbation or targeting lipid rafts since UDCA has a cholesterol skeleton which has a high affinity to lipid rafts. PVA-U25 was taken up in the cell at pH 7.4 because it interacted easily with the cell membrane owing to its high hydrophobicity (Figure 7e). In contrast, the DLS measurement results showed that PVA-U25 formed aggregates of about 1 µm in size at pH 6.5, although no aggregates could be observed from the confocal images. This result suggests that PVA-U25 formed aggregates at a size of approximately 1 µm, and they would be taken up by cell. PVA-U25 also showed smaller aggregates than PVA-U15 at pH 6.5 in confocal images, suggesting that PVA-U25 may prefer polymer-cell membrane to polymer-polymer interaction at 37 °C due to high G.D. of UDCA which allow easy insertion to cell membrane. In conclusion, we found that surface charge and hydrophilic-hydrophobic balance are important for regulating cell uptake and that high tumor microenvironment responsiveness can be achieved by designing polymers based on these factors.

**Figure 7.**
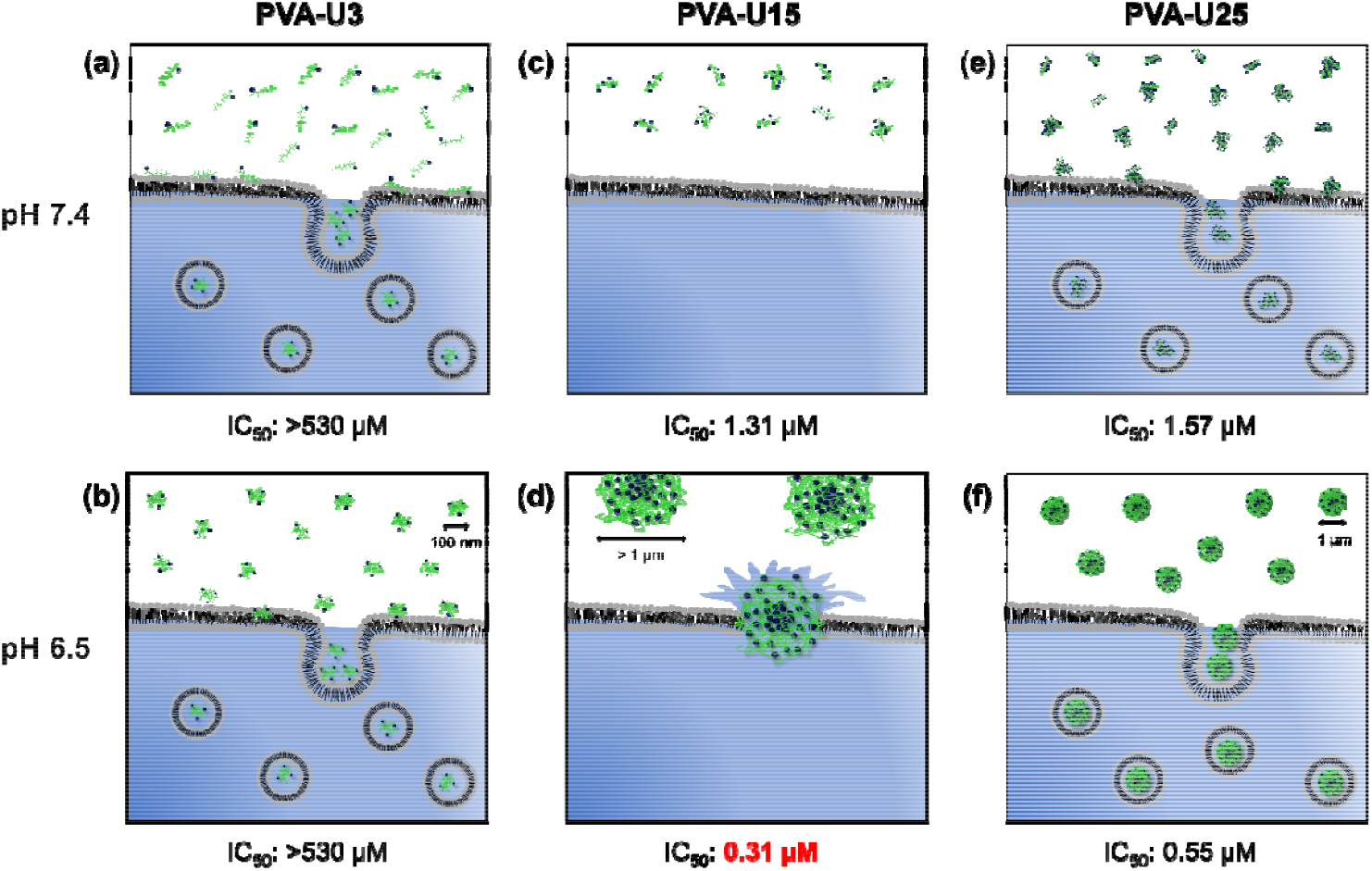
Schematic illustration of cell interaction of PVA-U3, PVA-U15, and PVA-U25 at pH 7.4 and pH 6.5. PVA-U3 was internalized in cell at pH 7.4 due to less anion charge, less hydrophobicity, and cellular insertion ability of slightly modified UDCA (a). In addition to these factors, size-dependent internalization was also possible at the nanoscale, which is invisible with confocal microscopy (approximately 100 nm) (b). PVA-U15 showed low cell membrane interaction at pH 7.4 (c) whereas it formed micro-scale aggregates (> 1 µm) pericellular without internalization in cell at pH 6.5 due to its suitable surface charge and hydrophilic-hydrophobic switching capacity (d). PVA-U25 was taken up into cell by high hydrophobicity at pH 7.4 (e). Although PVA-U25 was taken up by cells at a size scale that could not be observed with a confocal microscope, 1 *μ*m aggregates were formed at pH 6.5 in DLS, suggesting that PVA-U25 also showed aggregation at a size of 1 µm, resulting in cell internalization pH 6.5 (f).

We also investigated the cytotoxicity of PVA-U15 against not only MIAPaCa-2 cells but also other cancer cells and normal cells such as colon cancer (HT29), breast cancer (MDA-MB-231), lung cancer (A549), and NHDF. Each cell viability at pH 6.5 was lower than that at pH 7.4 (Figures S31a-31d) and IC_50_ values of PVA-U15 at pH 6.5 against each cell was lower than that at pH 7.4 (Figure S31e). The differences in IC_50_ values against HT29 cells, MDA-MB-231 cells, A549 cells, and NHDF between pH 7.4 and 6.5 were 2.2-fold, 3.2-fold, 6.0-fold, and 2.0-fold, respectively, suggesting the pH-responsive cytotoxicity of PVA-U15 varied by cell line although PVA-U15 universally showed tumor microenvironment responsive cytotoxicity. These results indicated that pH-responsive cytotoxicity in each cell line could be optimized by controlling the G.D. of PVA-U.

### 2.6. Antitumor efficacy and biocompatibility of PVA-U15

The antitumor efficacy of PVA-U15 was tested with a pancreatic cancer xenograft model. Nude mice subcutaneously inoculated with MIAPaCa-2cells were prepared and 100 µL of PBS solutions containing PVA, UDCA, and PVA-U15 were intratumorally injected (Figure 8a). Tumors in the mice treated with PVA grew from 81 mm^3^ at day 9 to 192 mm^3^ at day 21, an approximate doubling in size in two weeks. That of UDCA and PVA-U15 grew from approximately 80 mm^3^ to 144 mm^3^ and 101 mm^3^, respectively, suggesting that PVA-U15 significantly suppressed tumor growth compared to PVA (Figure 8b). Furthermore, the weight of mice treated with each solution stayed at around 20 g during administration, and no significant toxicity was observed in either treatment (Figure 8c). Therefore, it is suggested that PVA-U15 would have high antitumor efficacy and high biocompatibility based on its molecular design.

**Figure 8.**
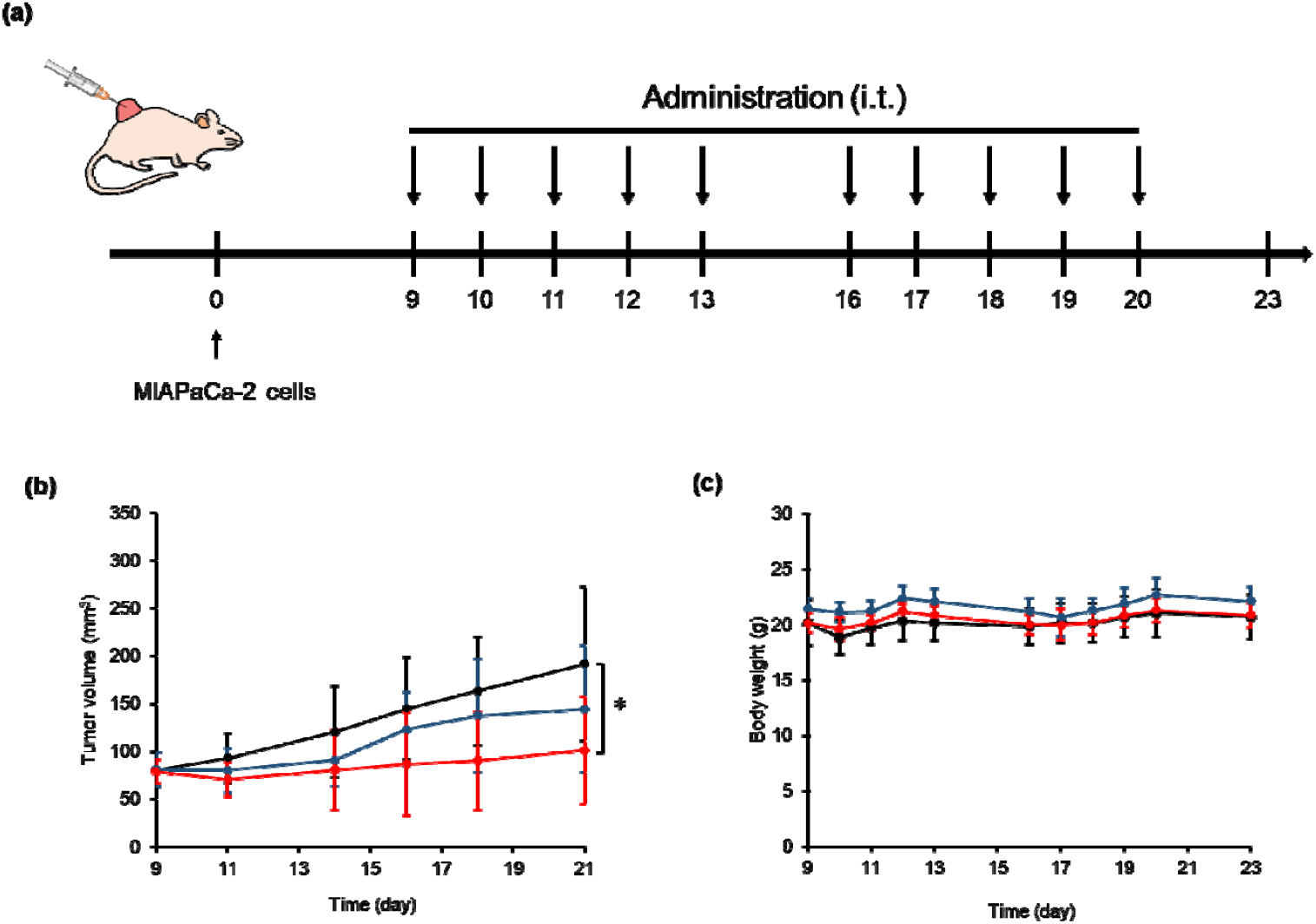
Antitumor efficacy of PVA-U15. (a) Treatment schedule for MIAPaCa-2 tumors. BALB/c nude mice were inoculated with MIAPaCa-2 cells on day 0. The tumor-bearing mice were randomized and treated with UDCA (144 µM), PVA (10 µg mL^−1^) or PVA-U15 (10 µg mL^−1^). 100 µL of each solution was intratumorally injected 5 days per week from day 9 to day 21. (b) Tumor growth in mice treated with PVA (black), UDCA (blue), and PVA-U15 (red). (c) Body weight of mice treated with PVA (black), UDCA (blue), and PVA-U15 (red). Statistical analysis was performed using Tukey test. Data are presented as mean ± S.D. (^*^*p* < 0.05).

## 3. Conclusion

In conclusion, we developed PVA-U where PVA was functionalized with UDCA that aggregated and induced cell death in response to a weak acidic condition. Although PVA-Us showed aggregation properties at pH 6.5, which is a tumor microenvironment, due to protonation of UDCA, the aggregation property, hydrophobicity, and surface charge were found to increase depending on the G.D. Notably, the difference in hydrophobicity of PVA-U15 between pH 7.4 and pH 6.5 was largest in PVA-Us, suggesting the hydrophilic-hydrophobic switching capacity could be optimized by tuning G.D. PVA-U15 showed the greatest difference in IC_50_ values between pH 7.4 and pH 6.5 against MIAPaCa-2 cells, suggesting that the tumor microenvironment selective cytotoxicity could be also optimized by controlling the G.D. This result was due to lower cellular uptake of PVA-U15 than PVA-U3 or PVA-U25, suggesting that PVA-U15 aggregated around the cell membrane in response to weak acid, thereby increasing the local concentration of UDCA and efficiently inducing cell membrane disruption. In addition, PVA-U3 would be easy to internalize in the cell due to low negative charge and hydrophobicity, and PVA-U25 would be taken up by the cell due to the preferential polymer-cell membrane interaction by the large amount of UDCA in the molecule. These results revealed the importance of molecular design to control the aggregation size, hydrophobicity, surface charge, and interaction with the cell membrane for efficiently inducing cancer cell death. Taken together, this study is expected to be important as a design direction for efficient induction of cell death in DDS studies with tumor-microenvironment responsive or membrane disrupting functions.

## 4. Experimental Section/Methods

### 4.1. Materials

Ursodeoxycholic acid (UDCA), methanesulfonyl chloride, dehydrated pyridine, 5 M hydrochloric acid (HCl), 5 M sodium hydroxide (NaOH), ethyl acetate, toluene, dimethyl sulfoxide (DMSO), and poly(vinyl) alcohol (PVA) with degree of polymerization 900-1,100 which was completely hydrolyzed, were purchased from Wako (Kyoto, Japan). WST-8 viable cell counting reagent and phosphate-buffered saline (PBS) were purchased from Nacalai Tesque (Kyoto, Japan). Methanol and sodium chloride (NaCl) were purchased from Kishida Chemical Co. Ltd. (Osaka, Japan). Rhodamine B isothiocyanate (RBITC), chloroform-*d*, and DMSO-*d*_*6*_ were purchased from Sigma-Aldrich (St. Louis, USA). CellTracker™ Deep Red dye and LysoTracker^TM^ deep red dye was purchased from Thermo Fisher Scientific (Massachusetts, USA). Cellulose tube was purchased from Japan Medical Science Co. Ltd. (Osaka, Japan).

### 4.2. Cell culture

MIAPaCa-2 cells, HT-29 cells, and A549 cells were purchased from the American Type Culture Collection (Manassas, VA, USA). MDA-MB-231 cells were purchased from KAC Co. Ltd. (Kyoto, Japan). Normal human dermal fibroblasts (NHDF) were purchased from Lonza (Basel, Switzerland) and maintained in Dulbecco’s modified Eagle medium (DMEM, Nacalai Tesque, Kyoto, Japan) supplemented with 10% fetal bovine serum (FBS, Thermo Fisher Scientific, Massachusetts, USA) and 1% antibiotic-antimycotic mixed stock solution (Nacalai Tesque, Kyoto, Japan) and non-essential amino acid (Thermo Fisher Scientific, Massachusetts, USA). Cells were cultured in 5% CO_2_ and 95% humidified air at 37 °C and passaged every 3 days.

### 4.3. Preparation of medium at a weak acidic pH

The medium at a weak acidic pH was prepared according to a previously reported method.^20^ 3% 1 mM HCl solution was added to the volume in the medium and incubated for 30 min to allow for equilibrium between the medium and CO_2_, then several µL of 1 mM HCl and 1 mM NaOH solution were added every 30 min until the pH of the medium was weakly acidic.

### 4.4. Mesylation of UDCA

The mesylation and modification of UDCA to PVA are depicted in Scheme S1. Mesylation of UDCA was performed using previously reported methods.^17,21^ Briefly, 2.95 g of UDCA (**1**) (7.50 mmol, 1.0 eq) was mixed with 50 mL of dehydrated pyridine and stirred at 0 °C in a nitrogen atmosphere. 700 µL of methanesulfonyl chloride (9.00 mmol, 1.2 eq) was added dropwise and the solution was stirred at 0 °C for 30 min. The reaction mixture was warmed to room temperature and then stirred for an additional 3 h. 100 mL of 5 M HCl was added to this reaction mixture and the solution was extracted with 100 mL of ethyl acetate three times. The 300 mL of mesylated UDCA solution was vacuum concentrated to around 50 mL by rotary evaporator (EYELA, N-1300, Tokyo, Japan) and 50 mL of toluene was added in mixture. The solution was vacuum concentrated and dried *in vacuo* for 3 days. The product was analyzed by proton nuclear magnetic resonance (^1^H NMR, JNM-GSX 400, JEOL, Tokyo, Japan). Mesylated UDCA (**2**): Yield: 3.83 mg, Substitution ratio: 96.8%. ^1^H NMR (400 MHz, chloroform-*d*): δ = 4.70-4.54 (br. m, 1 H), 3.56 (s, 1 H), 2.99 (s, 3 H, CH_3_SO_3_), 2.38 (m, 1 H), 2.24 (m, 1 H), 2.16-1.03 (m, 24 H), 0.97-0.84 (m, 6 H), 0.66 (s, 3 H) ppm.

### 4.5. Modification of PVA-U

Modification of PVA-U followed previously reported methods.^17^ 0.553 and 1.11 g of mesylated UDCA (**2**) (1.14 and 2.27 mmol, 2.5 and 5.0 eq / −OH) and 20 mg of PVA (4.55 × 10^−4^ units, 1.0 eq / −OH) were mixed with 10 mL of dehydrated DMSO and stirred at 100 °C in a nitrogen atmosphere. 91.5 and 183 µL of dehydrated pyridine (1.14 and 2.27 mmol, 2.5 and 5 eq / −OH) was added dropwise and the solution was stirred at 100 °C for 3 days. 1.11 g of mesylated UDCA (**2**) (2.27 mmol, 10 eq / −OH) and 10 mg of PVA (2.27 × 10^−4^ units, 1.0 eq / −OH) were mixed with 10 mL of dehydrated DMSO and stirred at 100 °C in a nitrogen atmosphere. 183 µL of dehydrated pyridine (2.27 mmol, 10 eq / −OH) was added dropwise and the solution was again stirred at 100 °C for 3 days. These reaction mixtures were dialyzed in cellulose tubes against methanol, 10 mM NaOH, and water for 3 days, 8 h, and overnight respectively. The dialyzed mixtures were then freeze-dried for 4 days. The product was analyzed by ^1^H NMR.

PVA-U3 (**3**): Yield: 11.5 mg, G.D.: 2.61%. ^1^H NMR (400 MHz, DMSO-*d*_6_): δ = 4.93 (s, 1H), 4.76 (s, 1H), 4.59 (s, 1H), 4.00 (s, 1H), 3.70 (s, 1H), 3.34 (s, 2H, H_2_O), 2.32-0.84 (m, 28H), 0.60 (s, 1H) ppm.

PVA-U15 (**3**): Yield: 21.3 mg, G.D.: 14.6%. ^1^H NMR (400 MHz, DMSO-*d*_6_): δ = 4.93 (s, 1H), 4.78 (s, 1H), 4.60 (s, 1H), 4.00 (s, 1H), 3.72 (s, 1H), 3.35 (s, 2H, H_2_O), 2.02-0.83 (m, 28H), 0.60 (s, 1H) ppm.

PVA-U25 (**3**): Yield: 27.1 mg, G.D.: 25.1%. ^1^H NMR (400 MHz, DMSO-*d*_6_): δ = 4.94 (s, 1H), 4.78 (s, 1H), 4.60 (s, 1H), 4.00 (s, 1H), 3.72 (s, 1H), 3.34 (s, 2H, H_2_O), 2.09-0.79 (m, 28H), 0.61 (s, 1H) ppm.

### 4.6. Measurement of pH-sensitive aggregation property of PVA-U

The pH-dependent light scattering intensity and size distribution were evaluated by dynamic light scattering (DLS) measurement. PVA (PVA-U0), PVA-U3, PVA-U15, and PVA-U25 were dissolved in DMSO to prepare each polymer solution at a concentration of 2 mg mL^−1^. 2.5 µL of these solutions were dissolved in 500 µL of PBS at pH 7.4, 7.2, 7.0, 6.8, 6.6, 6.5, 6.4, 6.2, 6.0, 5.8, and 5.6, respectively, to prepare polymer solutions containing 0.5% DMSO at a concentration of 10 µg mL^−1^. 500 µL of each solution was used for DLS measurements (Zetasizer Nano-ZS, Malvern, UK) 2 min after preparation.

### 4.7. Titration of UDCA and PVA-U

A p*K*a value was determined by titration. UDCA was dissolved in 5 M NaOH solution with 150 mM NaCl to prepare a UDCA solution at concentrations of 1, 5, 10, and 15 mM. PVA-U3, PVA-U15, and PVA-U25 were also dissolved in 5 M NaOH solution with 150 mM NaCl to prepare a PVA-U solution at concentrations of 100 µg mL^−1^, respectively. pH titration was carried out by adding small volumes (about 8 µL increment per sec) of 0.01 M HCl solution under stirring. The pH decreases in the range of 12 to 2 were measured by a titration system (COM-A19, HIRANUMA Co., Ltd., Ibaraki, Japan).

### 4.8. Critical aggregation concentration (CAC) of PVA-U

Light scattering intensity of DLS was used for CAC determination.^22^ PVA-U0, PVA-U3, PVA-U15, and PVA-U25 were dissolved in DMSO to prepare each polymer solution at concentrations of 0.02 ~ 60 mg mL^−1^. 2.5 µL of these solutions were dissolved in 500 µL PBS at pH 7.4 or 6.5, respectively, to prepare PVA-U solutions containing 0.5% DMSO at a concentration of 0.1 ~ 300 µg mL^−1^. 500 µL of each solution was used for DLS measurements (Zetasizer Nano-ZS, Malvern, UK) 2 min after preparation. The CAC value was determined by piecewise fitting model as follows,

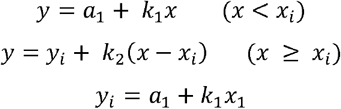

Where *y* is scattering intensity, *a*_l_ is the intercept, *x*_*i*_ is the intersection which is defined as CAC, and *y*_*i*_ is scattering intensity at a concentration of CAC.

### 4.9. Time-dependent size evaluation of PVA-U

The time-dependent mean size was evaluated by DLS measurement. PVA-U3, PVA-U15, and PVA-U25 were dissolved in DMSO to prepare each PVA-U solution at a concentrations of 2 mg mL^−1^. 2.5 µL of these solutions were dissolved in 500 µL of PBS at pH 7.4 and 6.5, respectively, to prepare polymer solutions containing 0.5% DMSO at a concentration of 10 µg mL^−1^. 500 µL of each solution was used for DLS measurements 2 min after preparation.

### 4.10. Zeta potential evaluation of PVA-U

Zeta potential was evaluated by DLS measurement. PVA-U0, PVA-U3, PVA-U15, and PVA-U25 were dissolved in DMSO to prepare each PVA-U solution at a concentrations of 10 mg mL^−1^. 2.5 µL of these solutions were dissolved in 500 µL of PBS at pH 7.4 and 6.5, respectively, to prepare polymer solutions containing 0.5% DMSO at a concentration of 50 µg mL^−1^. 500 µL of each solution was used for DLS measurements 2 min after preparation.

### 4.11. Contact angle measurement of PVA-U

Hydrophobicity of PVA-U was evaluated by contact angle measurement. 2 mg of PVA-U3, PVA-U15, and PVA-U25 was dissolved in 200 µL of DMSO to prepare a solution at each concentration of 10 mg mL^−1^. These solutions were thinly coated onto glass slides and vacuum dried to prepare polymer surfaces. 1x PBS at pH 7.4 and 6.5 were dropped on each polymer surface, and the contact angle was measured using contact angle meters (FAMAS, Kyowa Interface Co. Ltd., Saitama, Japan).

### 4.12. Evaluation of morphology of PVA-U by transmission electron microscopy (TEM) observation

1 mg of PVA-3, PVA-U15, and PVA-U25 were dissolved in 100 µL of DMSO to prepare a solution at each concentration of 10 mg mL^−1^. 2.5 µL of these solutions were dissolved in 500 µL of PBS at pH 7.4 and 6.5, respectively, to prepare polymer solutions containing 0.5% DMSO at a concentration of 10 µg mL^−1^. After 6 h of incubation, these samples were dropped on a carbon-coated copper disk (Nissin-EM, Tokyo, Japan) and freeze-dried overnight. Negative staining was used with phosphotungstic acid (PTA) solution. 20 mg of PTA was dissolved in 1 mL of Milli-Q to prepare 2% (w/v) PTA solution and their pH were adjusted to 7.4 and 6.5, respectively. Each sample was stained with 2% PTA solution adjusted at each pH before observation using a TEM microscope (H-7500, Hitachi, Tokyo, Japan).

The size distribution and circularity were analyzed from these images by Image J software. Size distribution of each PVA-U aggregate was analyzed by measuring the diameter of 30 aggregates each considered as a perfect circle. The circularity of each PVA-U was analyzed using the following formula:

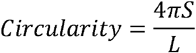

where S is area and L is the perimeter of each aggregate.

### 4.13. Concentration-dependent cytotoxicity of PVA-U

MIAPaCa-2 cells were seeded on a 96-well plate at 1.0 × 10^4^ cells per well for cell viability measurement and at 1, 2, 4, and 6 × 10^4^ cells per well for standard curve creation using a WST-8 assay. After 15 h incubation, cells for the standard curve were washed with 100 µL of PBS and incubated with 100 µL of 10% WST-8 viable cell counting reagent diluted in phenol red-free DMEM (Nacalai Tesque, Kyoto, Japan) with 1% antibiotics for 2 h at 37 °C and 5% CO_2_. 80 µL of the supernatant was transferred to another 96-well plate and the absorbance at 450 nm was measured using a plate reader. 0.9 mg of PVA-U0, PVA-U3, PVA-U15, and PVA-U25 were dissolved in 15 µL of DMSO to prepare each PVA-U solution at a concentration of 60 mg mL^−1^. These were diluted to 40, 20, 10, 2, 1, and 0.2 mg mL^−1^ with DMSO and 2.5 µL of these solutions were dissolved in 500 µL of DMEM supplemented with 10% FBS and 1% antibiotics at pH 7.4 and 6.5, respectively to prepare polymer solutions containing 0.5% DMSO at concentrations of 1, 5, 10, 50, 100, 200, and 300 µg mL^−1^. MIAPaCa-2 cells were treated with these media at 37 °C and 5% CO_2_ for 24 h. Phase contrast (Ph) microscopy images of MIAPaCa-2 cells incubated for 0.5 h and 24 h were taken using a Ph microscope. After 24 h, the absorbance of WST-8 was measured using the same procedure as for calibration of the standard curve. Cell viability was determined using the percentage of viable cells by calculating the number of cells in each sample using a standard curve and then using the following formula:

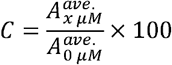

where C is the viable cell rate, 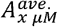 is the average number of living cells cultured with medium containing bile acid *x* µM for 24 h, and 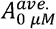 is that of the medium without bile acid. The fitting curve equation used to determine the IC_50_ value is as follows:

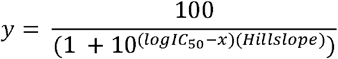

where *y* is the viable cell rate, *x* is the concentration of bile acid and Hill slope, which quantifies the steepness of the fitting curve, is calculated by GraphPad Prism.

### 4.14. Time dependent cytotoxicity of PVA-U15

MIAPaCa-2 cells were seeded on a 96-well plate at 1.0 × 10^4^ cells per well for cell viability measurement and at 1, 2, 4, and 6 × 10^4^ cells per well for standard curve creation using a WST-8 assay. After 15 h incubation, cells for the standard curve were washed with 100 µL of PBS and incubated with 100 µL of 10% WST-8 viable cell counting reagent diluted in phenol red-free DMEM (Nacalai Tesque, Kyoto, Japan) with 1% antibiotics for 2 h at 37 °C and 5% CO_2_. 80 µL of the supernatant was transferred to another 96-well plate and the absorbance at 450 nm was measured using a plate reader. 1.0 mg of PVA-U15 were dissolved in 100 µL of DMSO to prepare each PVA-U15 solution at concentrations of 10 mg mL^−1^. 2.5 µL of polymer solutions were dissolved in 500 µL of DMEM supplemented with 10% FBS and 1% antibiotics at pH 7.4 and 6.5, respectively to prepare polymer solutions containing 0.5% DMSO at a concentration of 10 µg mL^−1^. MIAPaCa-2 cells were treated with these media at 37 °C and 5% CO_2_ for 24 h. After incubation for 0.5 h, 4 h, 12 h, and 24 h, the absorbance of WST-8 was measured using the same procedure as for calibration of the standard curve.

### 4.15. Evaluation of cell membrane disruption of PVA-U

MIAPaCa-2 cells were seeded on a 96-well plate at 1.0 × 10^4^ cells for positive control, negative control, and treatment with each PVA-U. 0.9 mg of PVA-U0, PVA-U3, PVA-U15, and PVA-U25 were dissolved in 15 µL of DMSO to prepare each PVA-U solution at a concentration of 60 mg mL^−1^. These were diluted to 20 mg mL^−1^ with DMSO and 2.5 µL of these solutions were dissolved in 500 µL of DMEM supplemented with 10% FBS and 1% antibiotics at pH 7.4 and 6.5, respectively to prepare polymer solutions containing 0.5% DMSO at concentrations of 100 µg mL^−1^. MIAPaCa-2 cells 24 h after seeding were treated with these media at 37 °C and 5% CO_2_ for 24 h. After 24 h, 20 µL of lysis buffer and 5 µL of 100 mM NaOH were added to positive control and the samples at pH 6.5, respectively, to adjust the pH of all samples. DMEM was added to other samples to adjust the total volume of samples. 100 µL of supernatant of each sample were used for LDH assay (DOJINDO, Kumamoto, Japan) and absorbance of each supernatant was measured by plate reader. LDH release percentage was determined using the following formula:

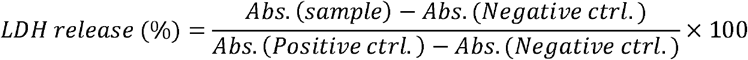

where *Abs, (sample)* is the absorbance of each sample, *Abs. (Positive ctrl*.) is the absorbance of positive control treated with lysis buffer, *Abs. (Negative ctrl.)* is the absorbance of negative control treated with DMEM.

### 4.16. Concentration-dependent cytotoxicity of PVA-U15 to various cancer cells and normal cells

Colon cancer (HT-29) cells, breast cancer (MDA-MB-231) cells, lung cancer (A549) cells, and NHDF were used for cytotoxicity evaluation of PVA-U15. Each cell type was seeded on a 96-well plate at 1.0 × 10^4^ cells per well for cell viability measurement and at 1, 2, 4, and 6 × 10^4^ cells per well for standard curve creation using a WST-8 assay. After 15 h incubation, cells for the standard curve were washed with 100 µL of PBS and incubated with 100 µL of 10% WST-8 viable cell counting reagent diluted in phenol red-free DMEM (Nacalai Tesque, Kyoto, Japan) with 1% antibiotics for 2 h at 37 °C and 5% CO_2_. 80 µL of the supernatant was transferred to another 96-well plate and the absorbance at 450 nm was measured using a plate reader. 0.9 mg of PVA-U0, PVA-U3, PVA-U15, and PVA-U25 were dissolved in 15 µL of DMSO to prepare each PVA-U solution at a concentration of 60 mg mL^−1^. These were diluted to 40, 20, 10, 2, 1, and 0.2 mg mL^−1^ with DMSO and 2.5 µL of these solutions were dissolved in 500 µL of DMEM supplemented with 10% FBS and 1% antibiotics at pH 7.4 and 6.5, respectively to prepare polymer solutions containing 0.5% DMSO at concentrations of 1, 5, 10, 50, 100, 200, and 300 µg mL^−1^. Each cell was treated with these media at 37 °C and 5% CO_2_ for 24 h. After 24 h, the absorbance of WST-8 was measured using the same procedure as for calibration of the standard curve. Cell viability was determined using the percentage of viable cells by calculating the number of cells in each sample using a standard curve and then using the following formula:

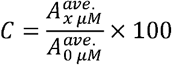

where C is the viable cell rate, 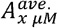 is the average number of living cells cultured with medium containing bile acid *x* µM for 24 h, and 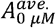 is that of the medium without bile acid. The fitting curve equation used to determine the IC_50_ value is as follows:

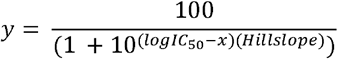

where *y* is the viable cell rate, *x* is the concentration of bile acid and Hill slope, which quantifies the steepness of the fitting curve, is calculated by GraphPad Prism.

### 4.17. Modification of rhodamine B isothiocyanate (RBITC)

The synthesis of PVA-U modified by RBITC (PVA-U-R) followed previously reported methods.^23^ 20 mg of PVA-U0 and 12.3 mg of RBITC were dissolved in 8 mL dehydrated DMSO. 5 mg PVA-U3, PVA-U15, and PVA-U25 (1.14 × 10^−4^ units, 1.0 eq / −OH) and 3.05 mg of RBITC (5.68 µmol, 0.05 eq / −OH) were dissolved in 2 mL of dehydrated DMSO. A droplet of pyridine and dibutyltin (IV) dilaurate were added in each solution and stirred at 95 °C for 2 h in a nitrogen atmosphere. Each reaction mixture was dialyzed using cellulose tubes against methanol for 6 days and against water for 1 day. The dialyzed mixture was freeze-dried for 7 days. The G.D. of each PVA-U was calculated by fluorescence intensity of RBITC using a spectrofluorometer (Jasco, Spectrofluorometer FP-8500, Tokyo, Japan). This was set at an excitation wavelength of 550 nm, fluorescence bandwidth of 2.5 nm, acquisition interval of 0.5 nm, temperature of 25 °C, and fluorescence wavelength of 450 nm to 650 nm. PVA-U0-R: G.D.: 2.96%. PVA-U3-R: G.D.: 0.75%. PVA-U15-R: G.D.: 0.80%. PVA-U25-R: G.D.: 0.66%.

### 4.18. Evaluation of cell adsorption of PVA-U-R over time using confocal images

MIAPaCa-2 cells were seeded on a glass based 24-well plate at 1.0 × 10^5^ cells per well for cell adsorption measurement. 15 µg CellTracker™ Deep Red were dissolved in 2.15 µL DMSO to prepare 10 mM CellTracker™ solution. 1 µL of this solution was mixed with 1 mL phenol red-free DMEM, and used for staining. 1 day after seeding MIAPaCa-2 cells, they were stained with 1 mL of CellTracker™ solution for 30 min and incubated for 1 day after washing with 1x PBS 3 times. 1 mg of PVA-U0, PVA-U3, PVA-U15, PVA-U25, PVA-U0-R, PVA-U3-R, PVA-U15-R, and PVA-U25-R were dissolved in 100 µL of DMSO, respectively, to prepare polymer solutions at a concentration of 10 mg mL^−1^. These solutions were diluted 5-fold with DMSO. The amount of RBITC was adjusted by mixing 0.7, 2.7, 2.5, and 3.0 µL of PVA-U0-R, PVA-U3-R, PVA-U15-R, and PVA-U25-R solution with 4.3, 2.3, 2.5, and 2.0 µL of PVA-U3, PVA-U15, and PVA-U25 solutions, respectively. 5 µL of each polymer solution at a concentration of 2 mg mL^−1^ was dissolved in 1 mL of phenol red-free DMEM supplemented with 10% FBS and 1% antibiotics at pH 7.4 and 6.5, to prepare a medium containing each polymer at a concentration of 10 µg mL^−1^. MIAPaCa-2 cells were treated with these media at pH 7.4 and 6.5 under 37 °C and 5% CO_2_ using an ultrasmall chamber for cell culture (BLAST Co. Ltd., SV-140A, Kanagawa, Japan). MIAPaCa-2 cells were monitoring for 24 h with a confocal laser scanning microscope (FV3000, Olympus, Tokyo, Japan) and confocal images were taken every 1 h. Intracellular brightness intensity (B.I.) of PVA-U-R was quantified by enclosing cell contour from the CellTracker images. Pericellular B.I. was calculated by subtracting the intracellular B.I. from the B.I. of a 5 *μ*m enlarged treatment of the contour enclosing the cell. 15 cells were randomly selected for the analysis and each average value of B.I. was calculated on 11 samples excluding the top two and bottom two B.I.

### 4.19. Evaluation of endocytosis of PVA-U

MIAPaCa-2 cells were seeded on a glass based 24-well plate at 1.0 × 10^5^ cells per well for cell adsorption measurement. 1 mg of PVA-U0, PVA-U3, PVA-U15, PVA-U25, PVA-U0-R, PVA-U3-R, PVA-U15-R, and PVA-U25-R were dissolved in 100 µL of DMSO, respectively, to prepare polymer solutions at a concentration of 10 mg mL^−1^. These solutions were diluted 5-fold with DMSO. The amount of RBITC was adjusted by mixing 0.7, 2.7, 2.5, and 3.0 µL of PVA-U0-R, PVA-U3-R, PVA-U15-R, and PVA-U25-R solution with 4.3, 2.3, 2.5, and 2.0 µL of PVA-U3, PVA-U15, and PVA-U25 solutions, respectively. 5 µL of each polymer solution at a concentration of 2 mg mL^−1^ was dissolved in 1 mL of phenol red-free DMEM supplemented with 10% FBS and 1% antibiotics at pH 7.4 and 6.5, to prepare a medium containing each polymer at a concentration of 10 µg mL^−1^. MIAPaCa-2 cells were treated with these media at pH 7.4 and 6.5. After 4 h incubation, 200 µL lysotracker at a concentration of 50 nM was added to these samples for 30 min and washed with phenol red free DMEM 3 times. These samples were observed with a confocal laser scanning microscope and analyzed by line scanning the cells on the images.

### 4.20. Evaluation of endocytosis inhibition of the cell treated with PVA-U-R

MIAPaCa-2 cells were seeded on a glass-based 24-well plate at 1.0 × 10^5^ cells per well for cell adsorption measurement. 1 mg of PVA-U0, PVA-U3, PVA-U15, PVA-U25, PVA-U0-R, PVA-U3-R, PVA-U15-R, and PVA-U25-R were dissolved in 100 µL of DMSO, respectively, to prepare polymer solutions at a concentration of 10 mg mL^−1^. These solutions were diluted 5-fold with DMSO. The amount of RBITC was adjusted by mixing 0.7, 2.7, 2.5, and 3.0 µL of PVA-U0-R, PVA-U3-R, PVA-U15-R, and PVA-U25-R solution with 4.3, 2.3, 2.5, and 2.0 µL of PVA-U0, PVA-U3, PVA-U15, and PVA-U25 solutions, respectively. 5 µL of each polymer solution at a concentration of 2 mg mL^−1^ was dissolved in 1 mL of phenol red-free DMEM supplemented with 10% FBS and 1% antibiotics at pH 7.4 and 6.5, to prepare a medium containing each polymer at a concentration of 10 µg mL^−1^. MIAPaCa-2 cells were treated with these media at pH 7.4 and 6.5 under 4 °C in a refrigerator for 4 h and observed with a confocal laser scanning microscope. Intracellular B.I. of PVA-U-R was quantified by enclosing cell contour from the CellTracker images. 15 cells were randomly selected for the analysis and each average value of B.I. was calculated on 11 samples excluding the top two and bottom two B.I.

### 4.21. *In vivo* tumor treatment

All animal studies were performed in accordance with animal protocols approved by the Institutional Animal Care and Use Committee and institutional guidelines. Specific pathogen-free 5-week female BALB/c nude mice were purchased from The Jackson Laboratory Japan, Inc. (Yokohama, Japan). 5 × 10^6^ MIAPaCa-2 cells in 100 µL of Hank’s balanced salt solution were subcutaneously injected into each mouse dorsum. When tumor volumes reached approximately 100 mm^3^, the mice were randomly divided into 3 groups (8 mice in each group), and treated with the PVA-U0, PVA-U15 or UDCA as follows. 10 mg mL^−1^ PVA-U0 and PVA-U15, and 14.4 mM UDCA solution in DMSO were prepared and 10 µL of each solution was dissolved in 990 µL of PBS, to prepare the solution containing each polymer at a concentration of 100 µg mL^−1^ and UDCA at a concentration of 144 µM. 100 µL of each solution was intratumorally injected 5 days per week for 2 weeks, and tumor volume and body weight were recorded during the treatment. Tumor volume (mm^3^) was calculated as length × width^2^ × 0.5.

### 4.22. Statistical analysis

The results are expressed as mean ± standard deviation (S.D.). Student’s t-tests were used to test for differences between group means.

## Acknowledgements

This work was supported by a JST SPRING (JPMJSP2138) from Japan Science and Technology Agency (JST). This study was financially supported by KAKENHI JP22H05131, JP22H05138, JP22H05140, JP22H05141, JP25H01220, JP22K21348, and JP21H04634 from Japan Society for Promotion of Science (JSPS). This study was also supported by JPJSBP120239201, JPJSBP120252301, and Y2024L0906033 from JSPS, COI-NEXT (JPMJPF2009) from JST.

## Data Availability Statement

Received: ((will be filled in by the editorial staff))

Revised: ((will be filled in by the editorial staff))

Published online: ((will be filled in by the editorial staff))

## Supporting Information

Supporting Information is available from the Wiley Online Library or from the author.

