## Supplemental information for "Escape from Cell Uptake: Drug-Free Cancer Therapeutics Regulated Hydrophobicity and Negative Charge"

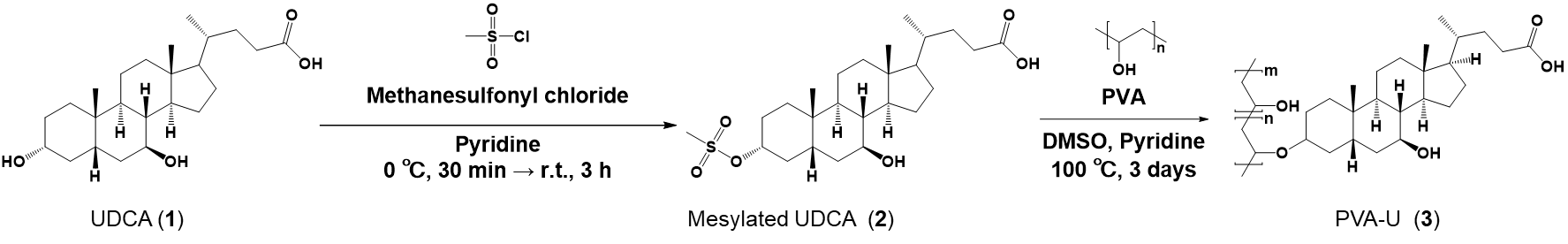

**Scheme S1.** Synthetic scheme of PVA-U.

**Table S1.** Equivalent ratio of UDCA-Ms, G.D., and yield of PVA-U.

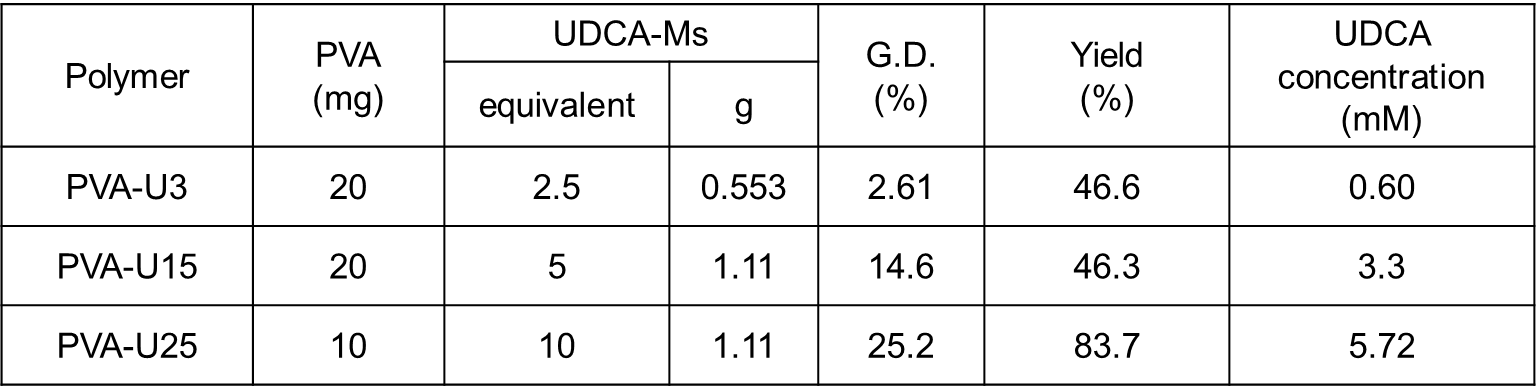

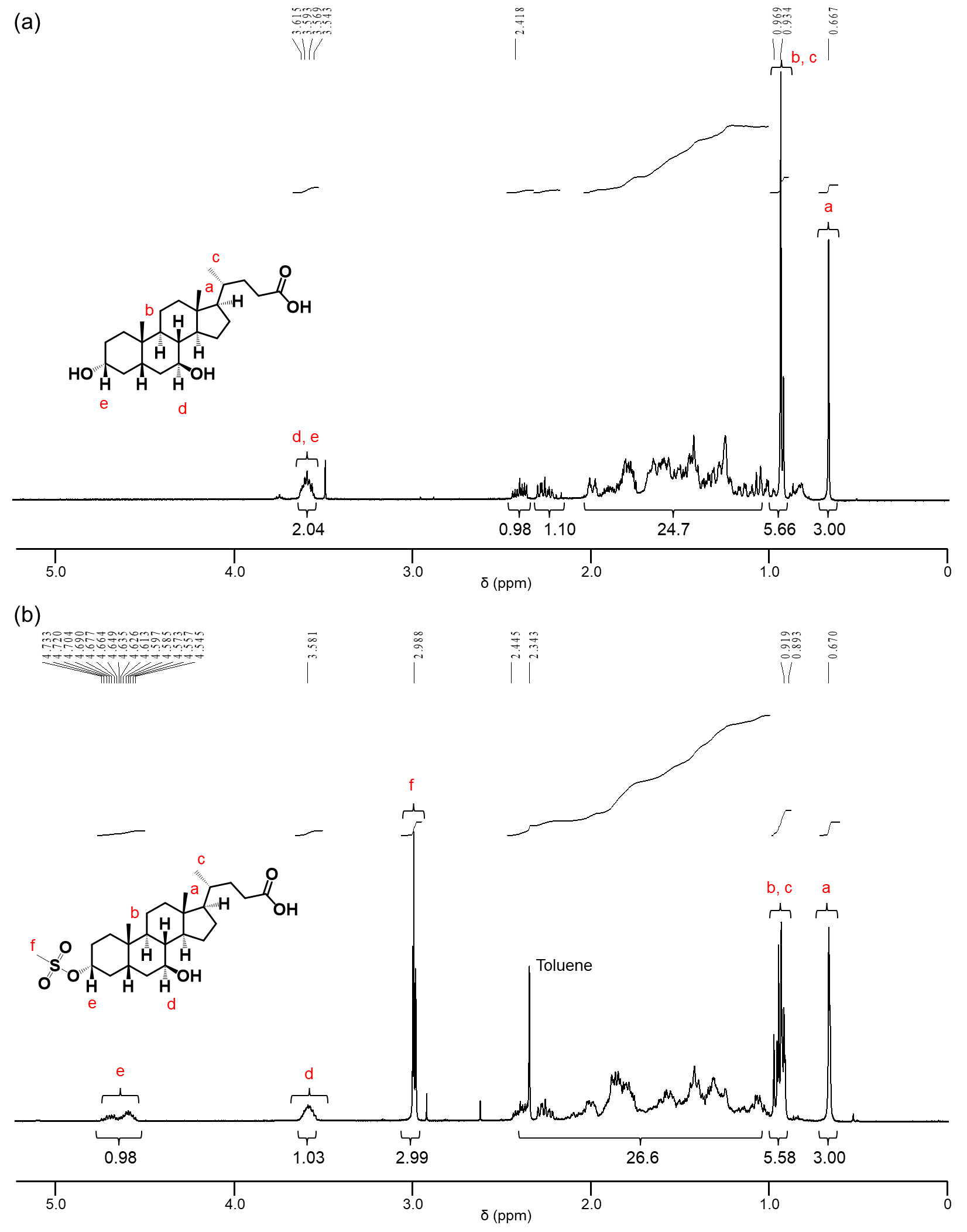

**Figure S1.** ^1^H NMR spectrum (Chloroform-*d* 400 MHz, 25 ˚C) of UDCA (a) and UDCA-Ms (b).

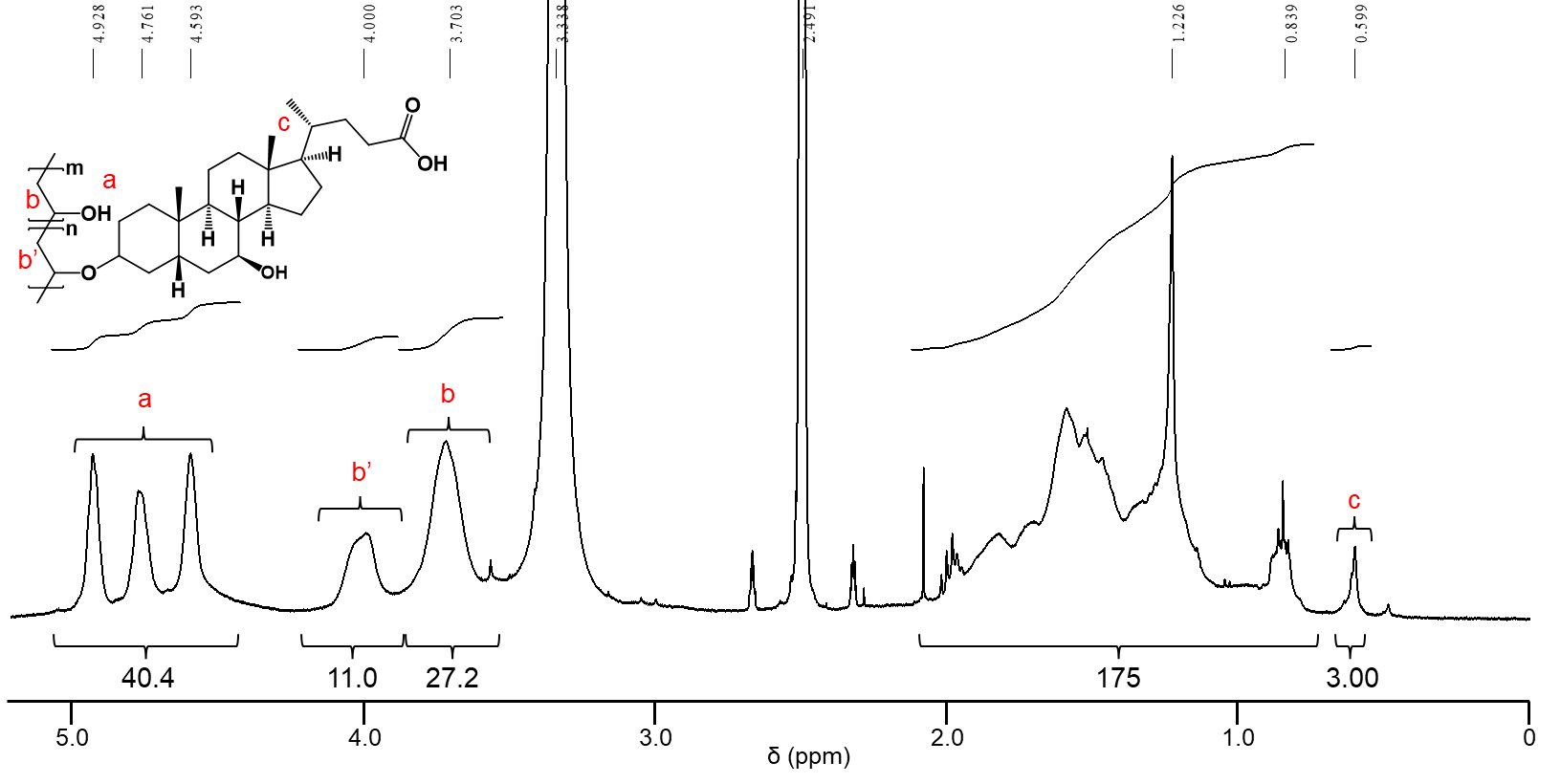

**Figure S2.** ^1^H-NMR spectrum (DMSO-*d_6_* 400 MHz, 25 ˚C) of PVA-U3.

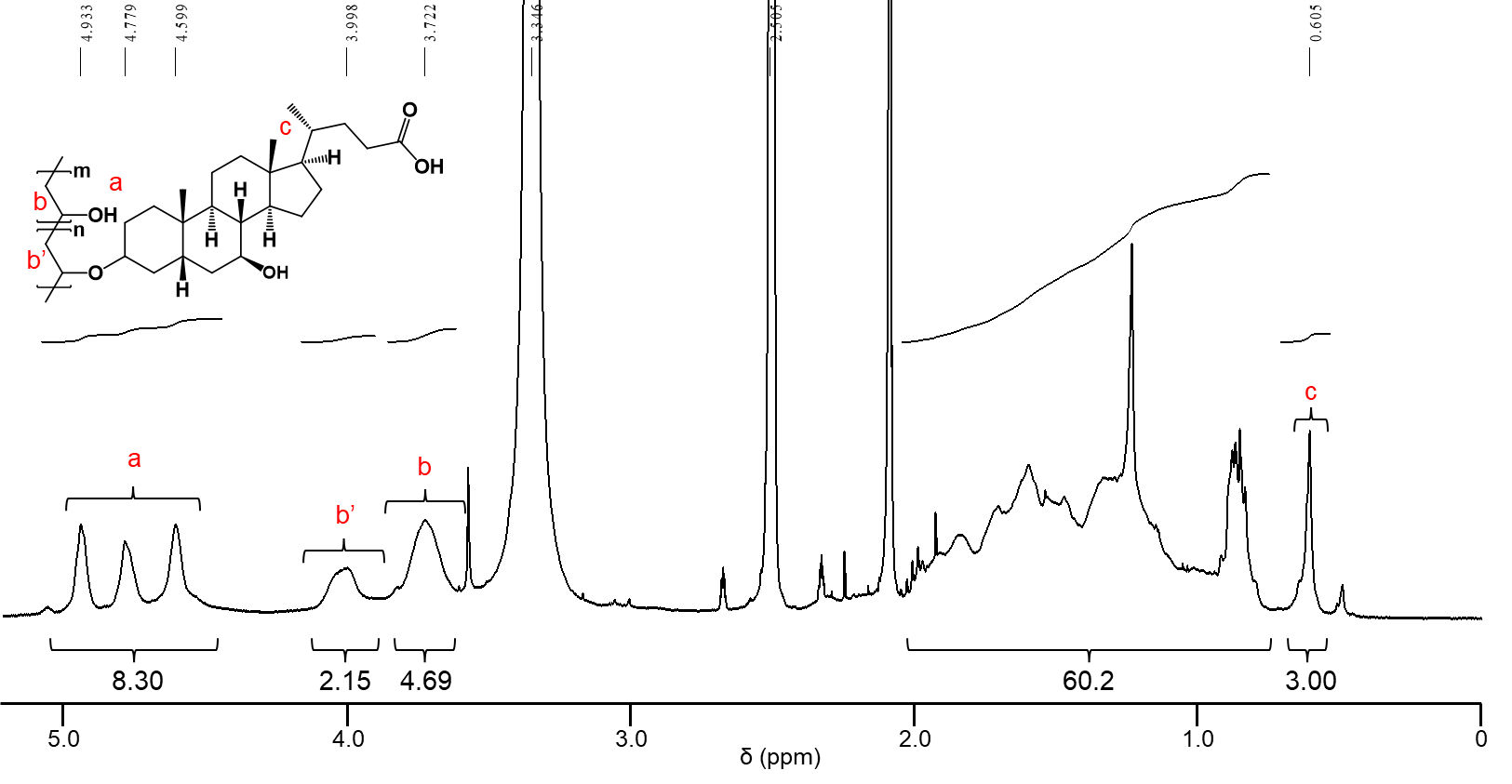

**Figure S3.** ^1^H-NMR spectrum (DMSO-*d_6_* 400 MHz, 25 ˚C) of PVA-U15.

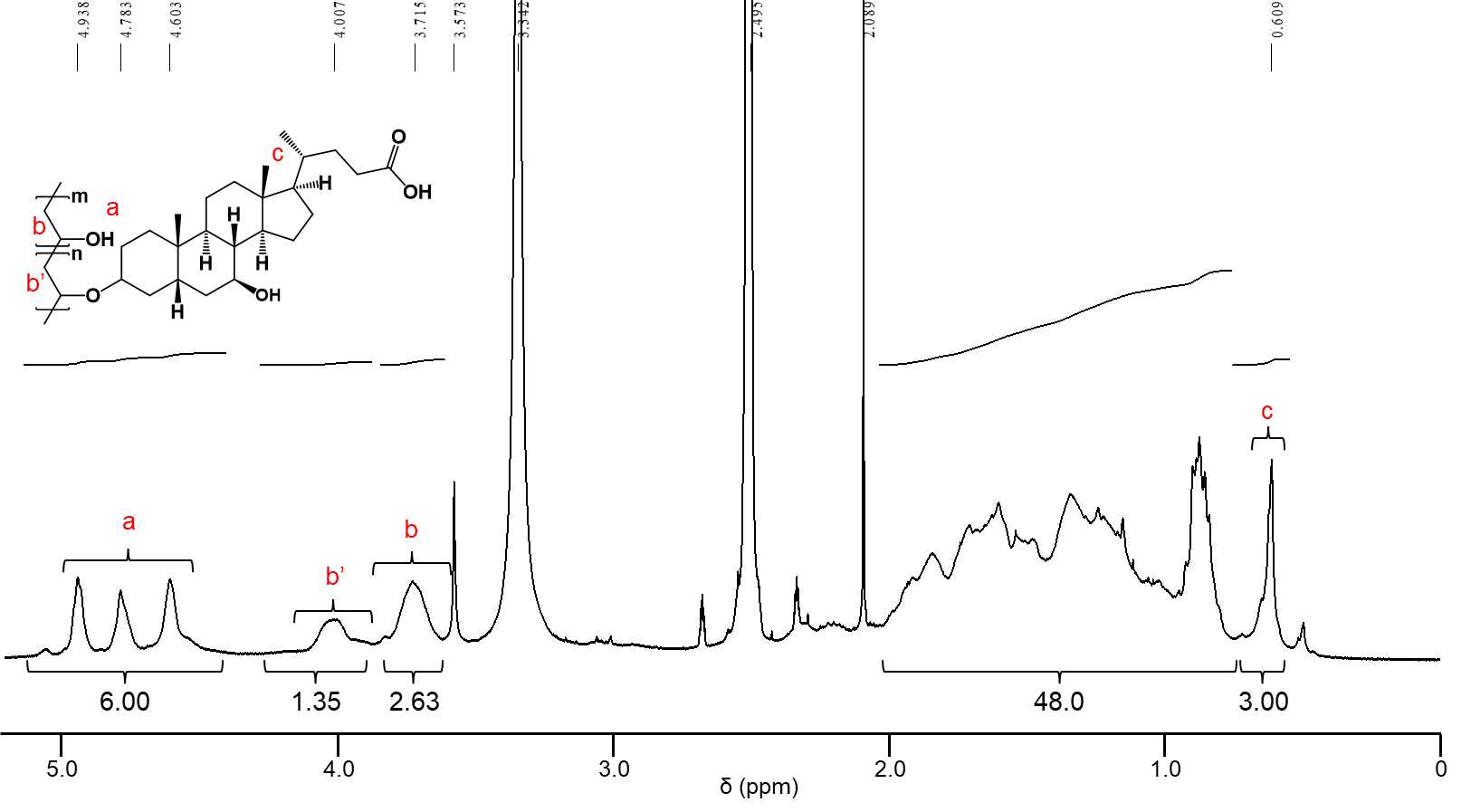

**Figure S4.** ^1^H-NMR spectrum (DMSO-*d_6_* 400 MHz, 25 ˚C) of PVA-U25.

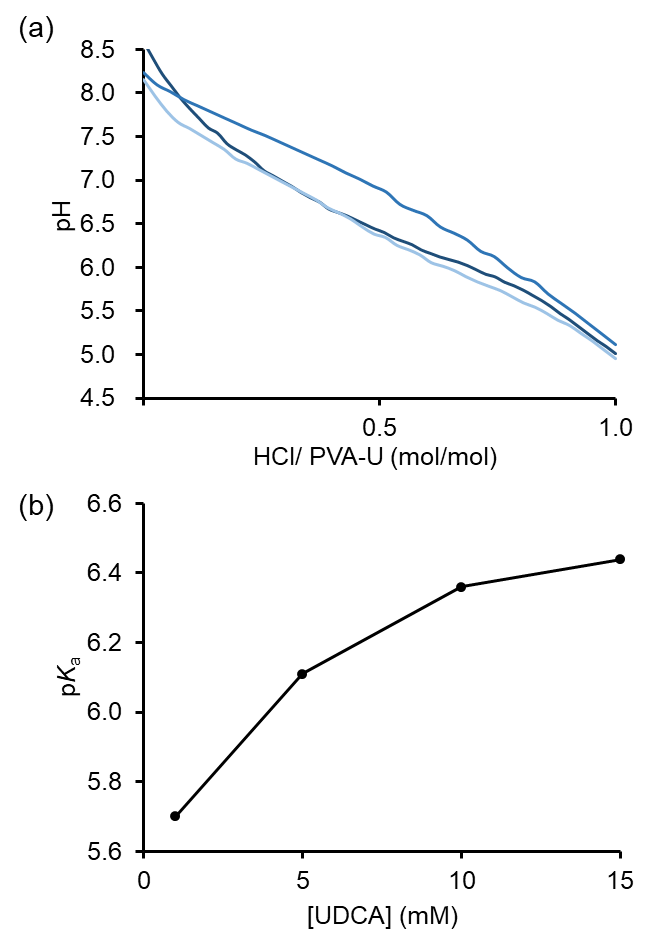

**Figure S5.** (a) Titration curve of PVA-U3 (light blue), 15 (blue), 25 (dark blue) at a concentration of 100 µg mL^-1^ in NaCl solution. (b) p*K*_a_ value of UDCA at each concentration.
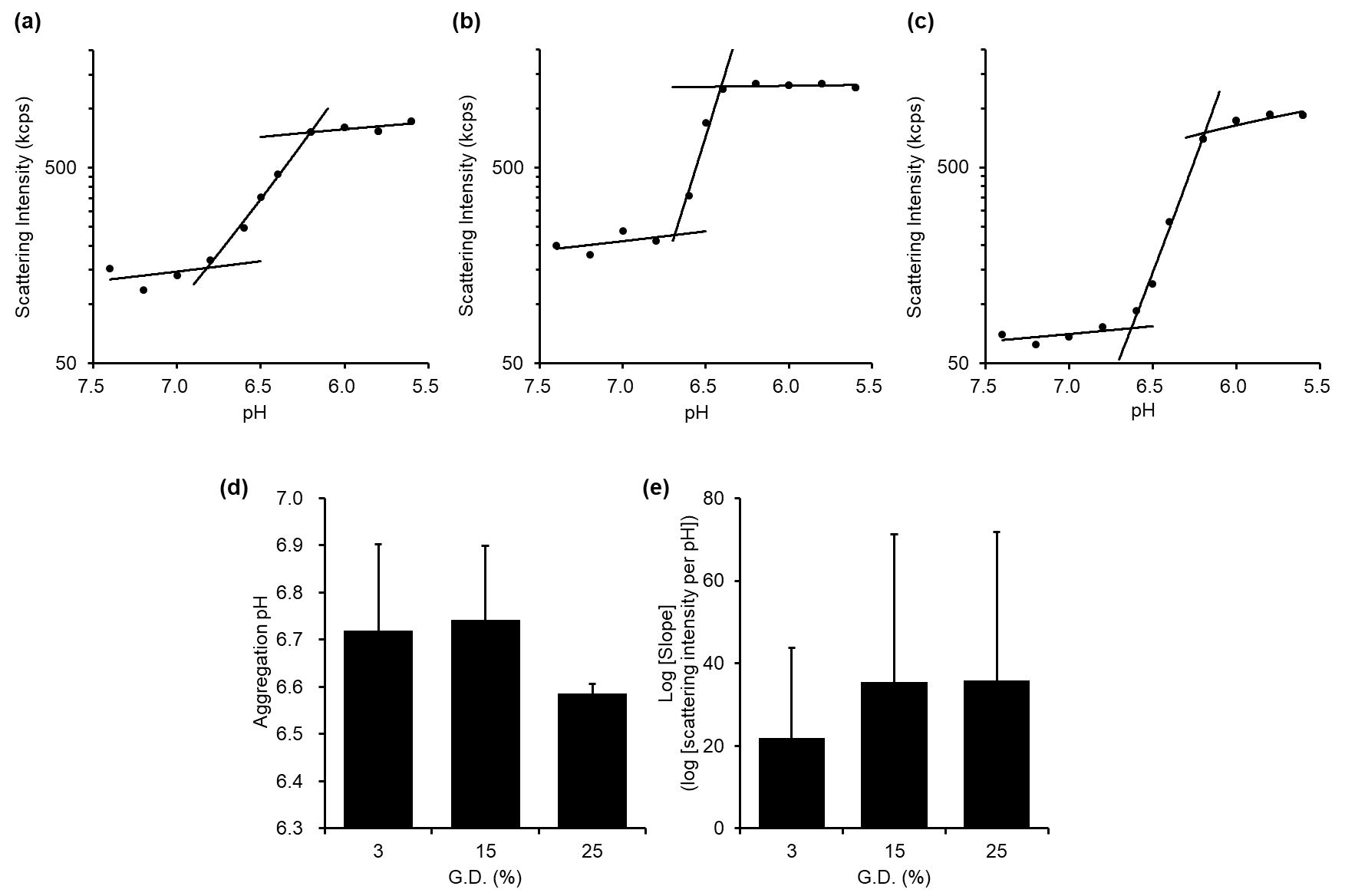

**Figure S6.** Representative fitting results of pH-dependent light scattering intensity of 10 µg mL^-1^ PVA-U3 (a), PVA-U15 (b), and PVA-U25 (c) in PBS immediately after preparation at 37 °C. (d) Aggregation pH of based on fitting curve in pH-dependent scattering intensity of PVA-U. (e) Logarithm slope of based on fitting curve in pH-dependent scattering intensity of PVA-U.

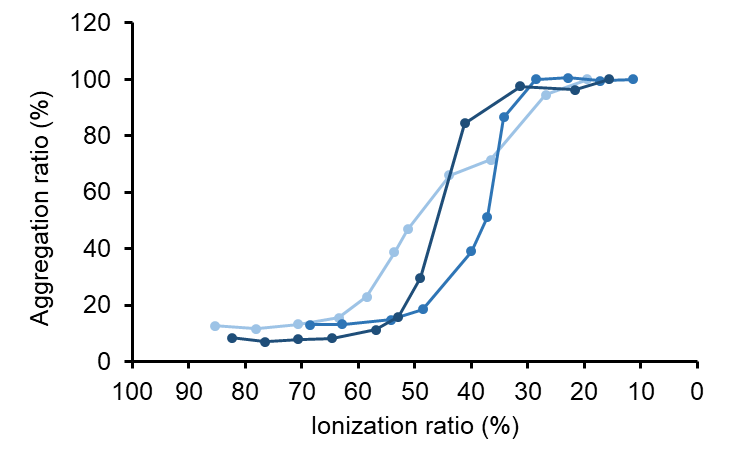

**Figure S7.** Aggregation ratio of PVA-U normalized by each scattering intensity at pH 5.6 against ionization ratio.

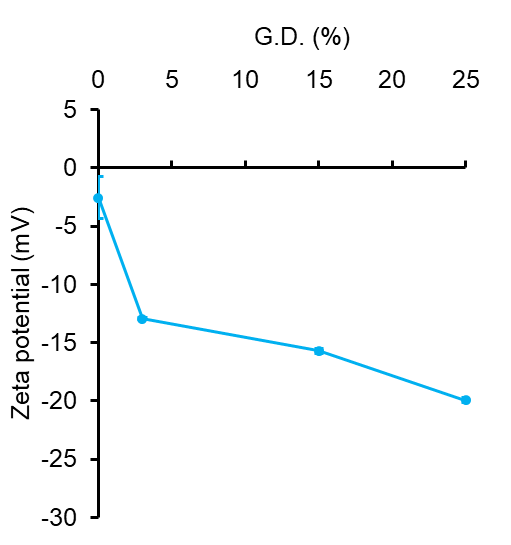

**Figure S8.** Zeta potential values of PVA-U at 5.6 (blue) immediately after preparation at 37 °C.

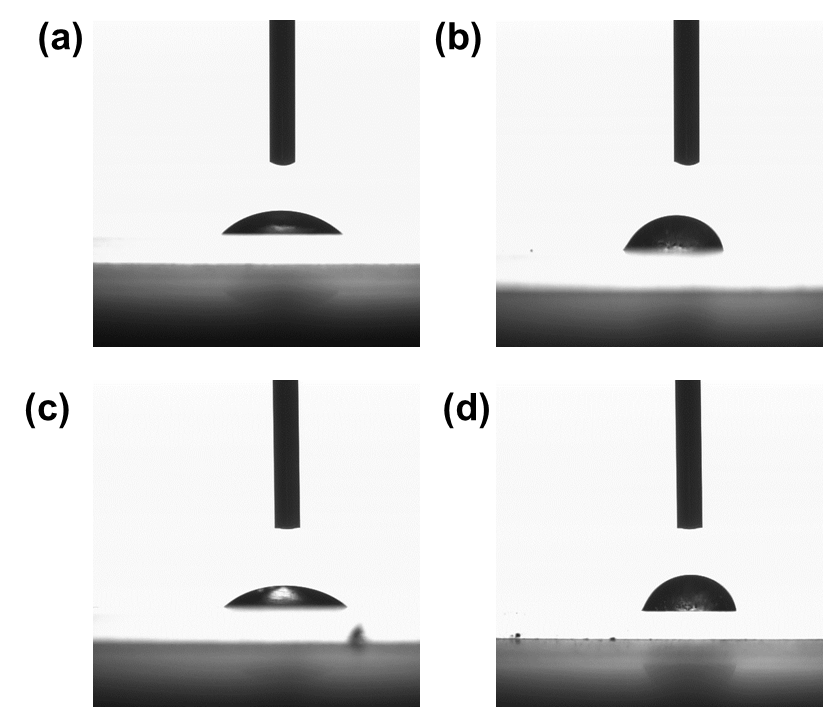

**Figure S9**. Images of droplet of 1X PBS at pH 7.4 (a, c) and 6.5 (b, d) to slide grass coated by PVA-U3 and PVA-U25.

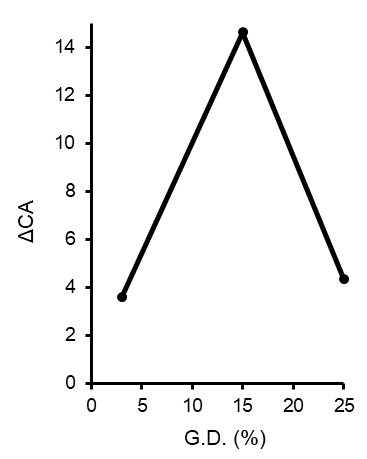

**Figure S10.** Contact angle difference of PVA-U3, PVA-U15, and PVA-U25

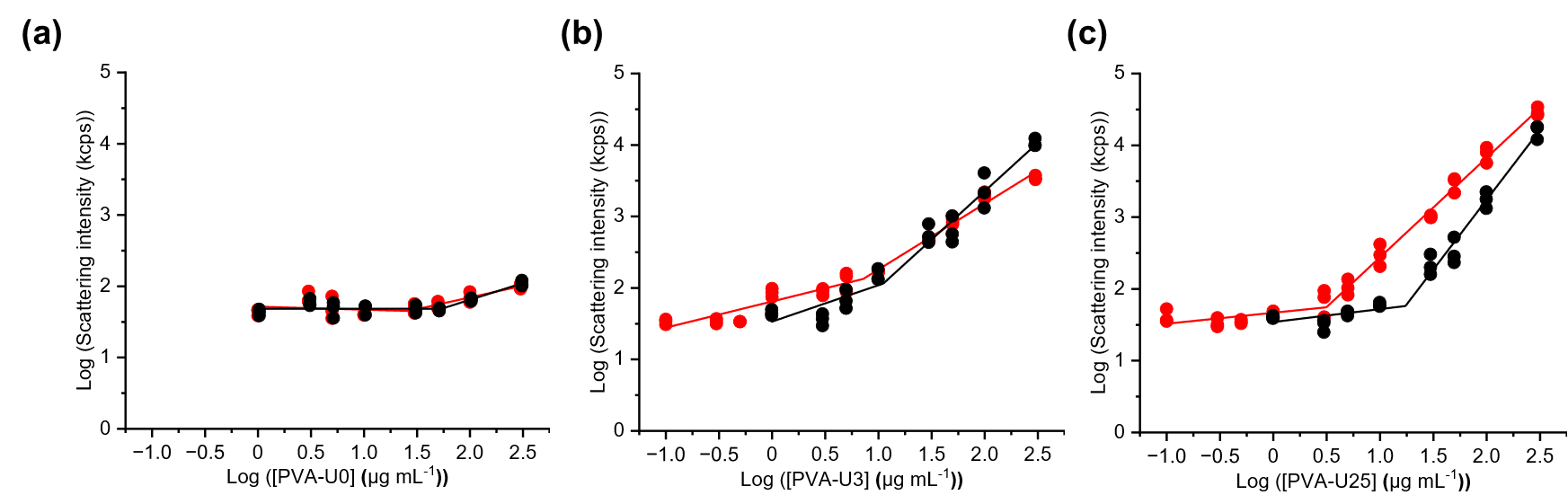

**Figure S11.** Concentration-dependent light scattering intensity of PVA-U0 (a), PVA-U3 (b), and PVA-U25 (c) in PBS at pH 7.4 (black) and pH 6.5 (red) immediately after preparation at 37 °C.

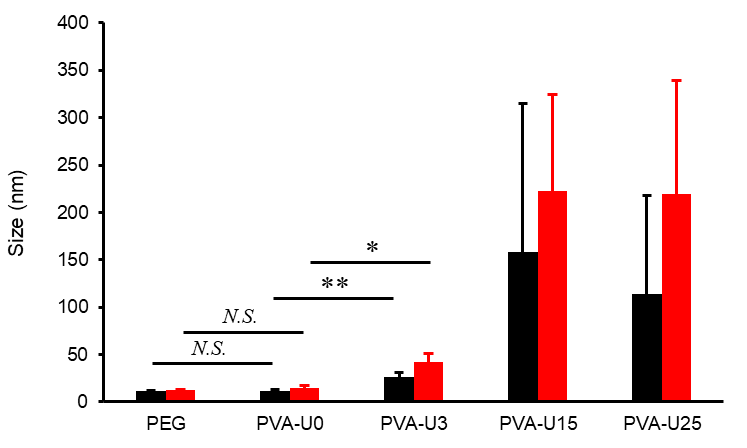

**Figure S12.** Mean size of PEG, PVA-U0, PVA-U3, PVA-U15, and PVA-U25 at pH 7.4 (black) and 6.5 (red) at a concentration of 300 µg mL^-1^. Statistical analysis was performed using unpaired two-tailed Student’s *t*-test. Data are presented as mean ± S.D. (*N.S.*: no significant difference, **p* < 0.05, ***p* < 0.01)

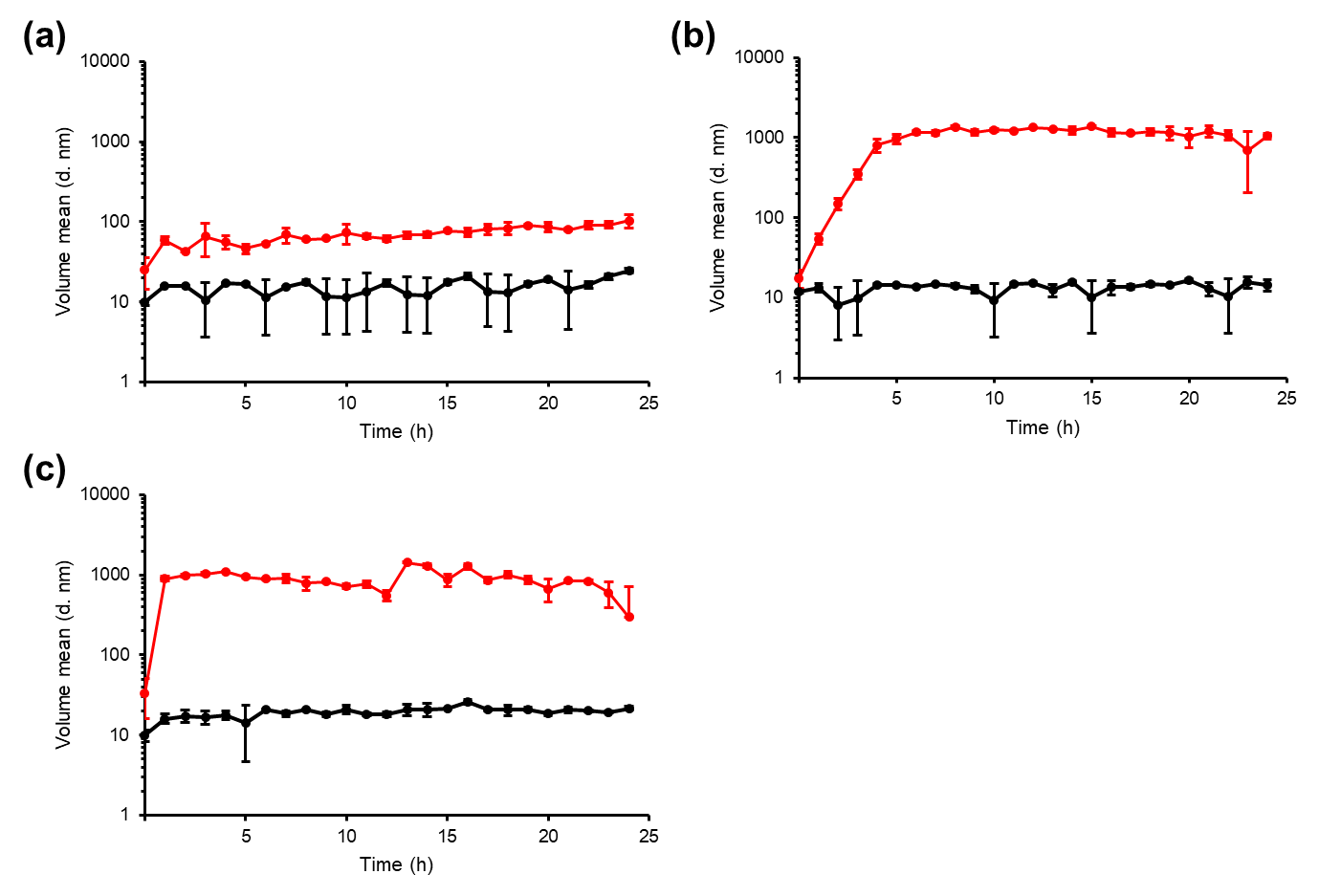

**Figure S13.** Time-dependent mean size of 10 µg mL^-1^ PVA-U3 (a), PVA-U15 (b), and PVA-U25 (c) at pH 7.4 (black) and 6.5 (red) for 24 hours incubation.

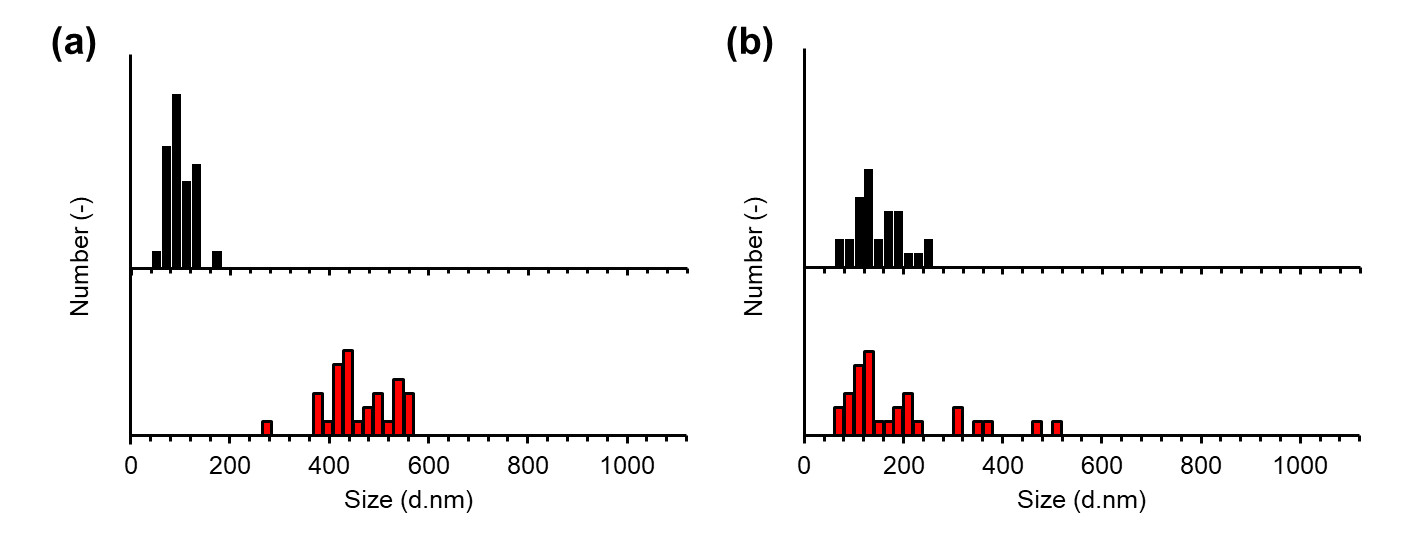

**Figure S14.** Size distribution of PVA-U3 (a) and PVA-U25 (b) aggregates at pH 7.4 (black) and pH 6.5 (red) analyzed from TEM images.

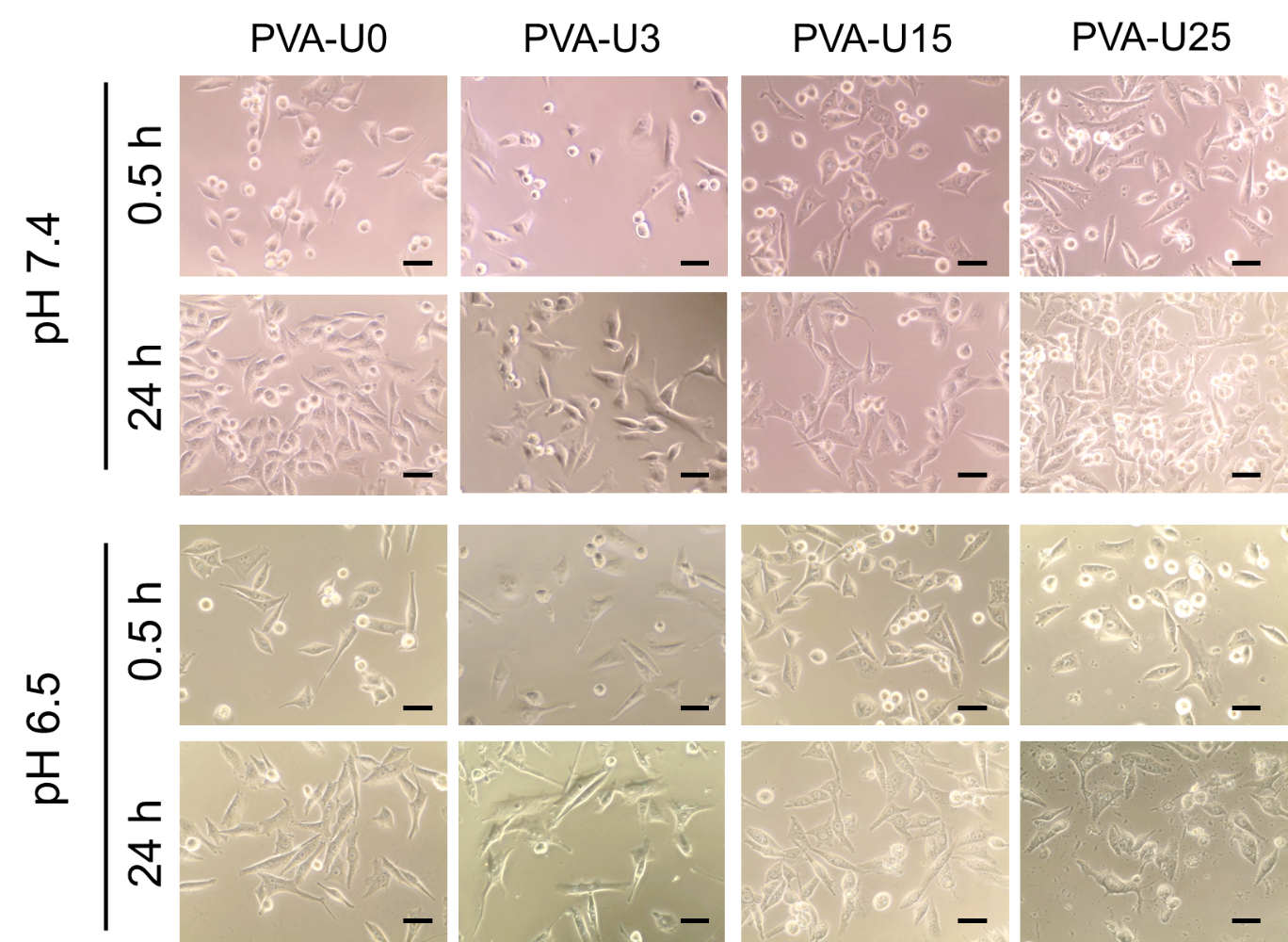

**Figure S15.** Phase contrast images of MiaPaCa-2 cells treated with 50 µg mL^-1^ PVA-U0, PVA-U3, PVA-U15, and PVA-U25 at pH 7.4 and pH 6.5 after 0.5 h and 24 h incubation. Scale bar is 50 µm.

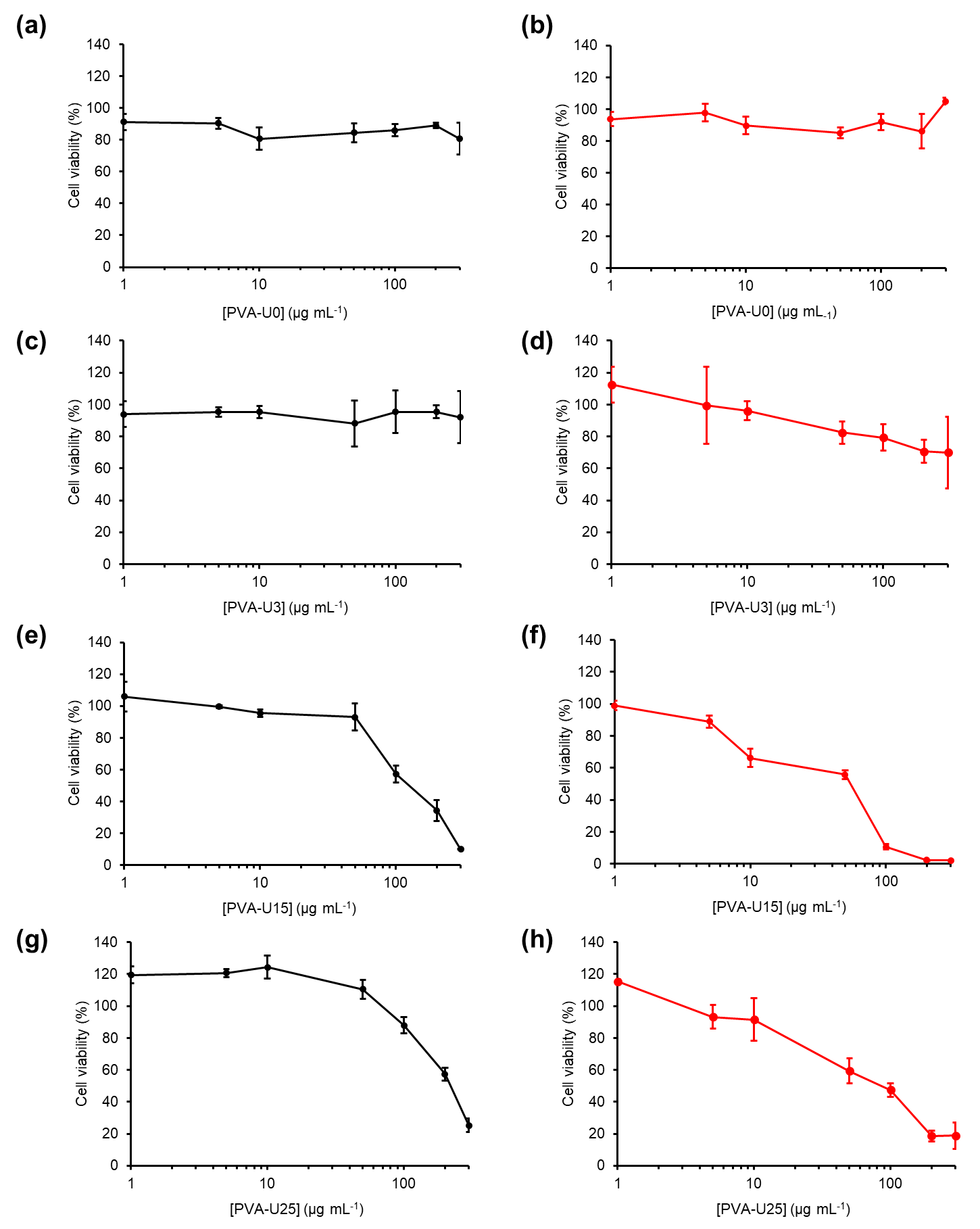

**Figure S16.** Cell viability of MiaPaCa-2 cells treated with PVA-U0 (a and b), PVA-U3 (c and d), PVA-U15 (e and f), and PVA-U25 (g and h) at pH 7.4 (black) and pH 6.5 (red) after 24 h incubation

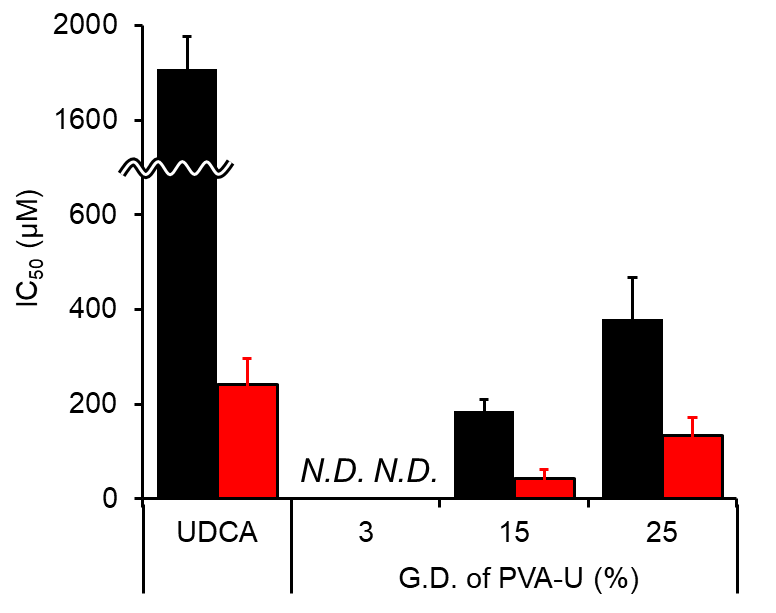

**Figure S17.** IC_50_ values of MiaPaCa-2 cells treated with UDCA, PVA-U3, PVA-U15, and PVA-U25 for 24 h incubation at pH 7.4 (black) and 6.5 (red) based on UDCA concentration. *N.D.*: not determined.

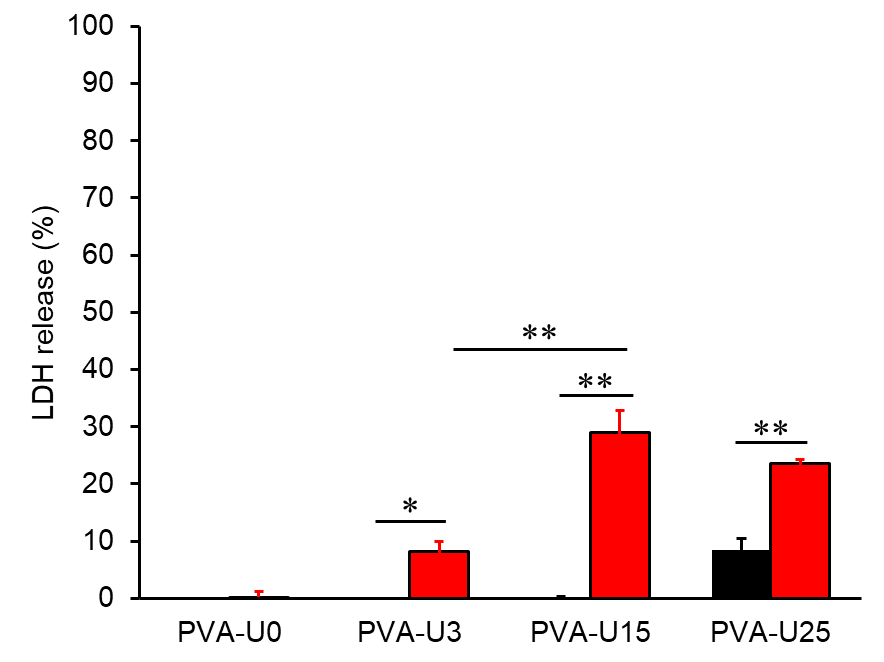

**Figure S18.** Leakage of LDH in MIAPaCa-2 cells treated with PVA-U0, PVA-U3, PVA-U15, and PVA-U25 at pH 7.4 (black) and 6.5 (red). Statistical analysis was performed using unpaired two-tailed Student’s *t*-test. Data are presented as mean ± S.D. (**p* < 0.05, ***p* < 0.01)

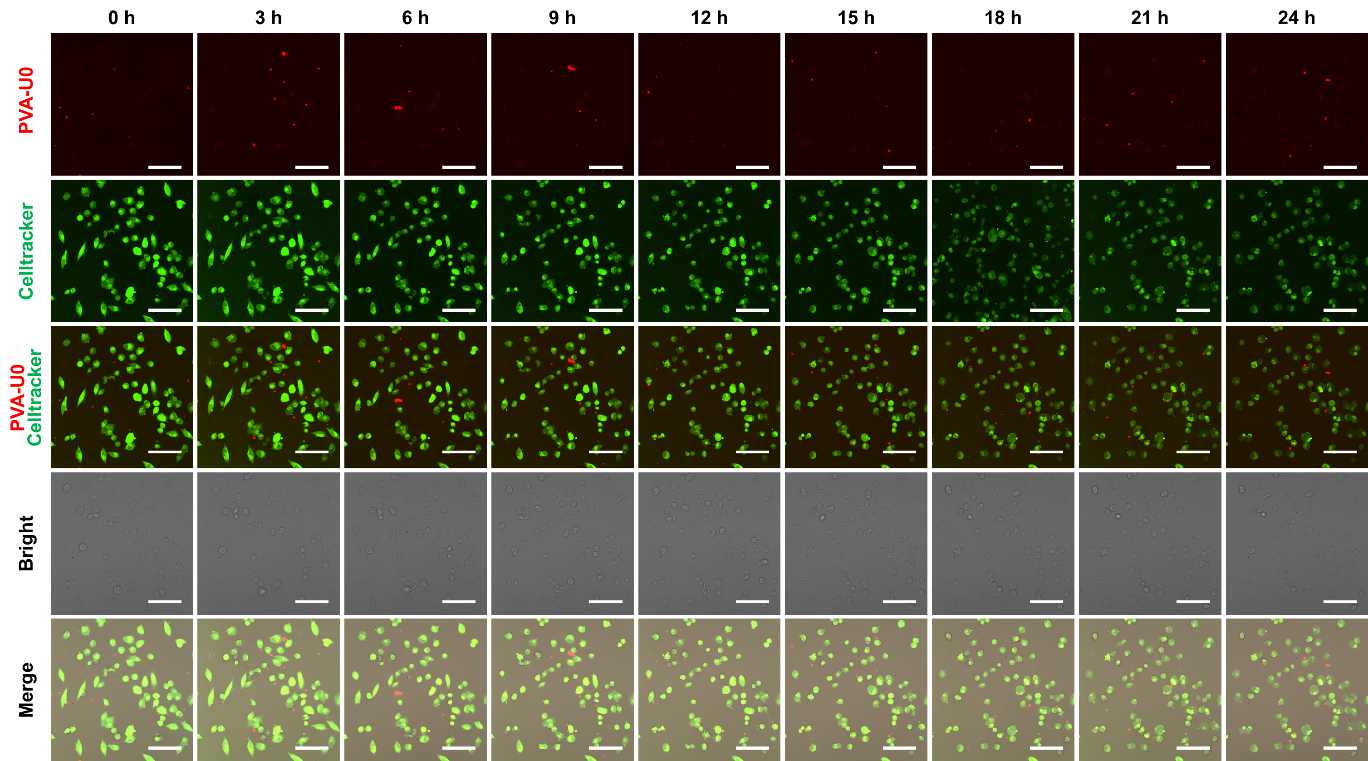

**Figure S19.** Confocal and bright filed images of MiaPaCa-2 cells treated with Celltracker and PVA-U0 at pH 7.4. Scale bar is 50 µm.

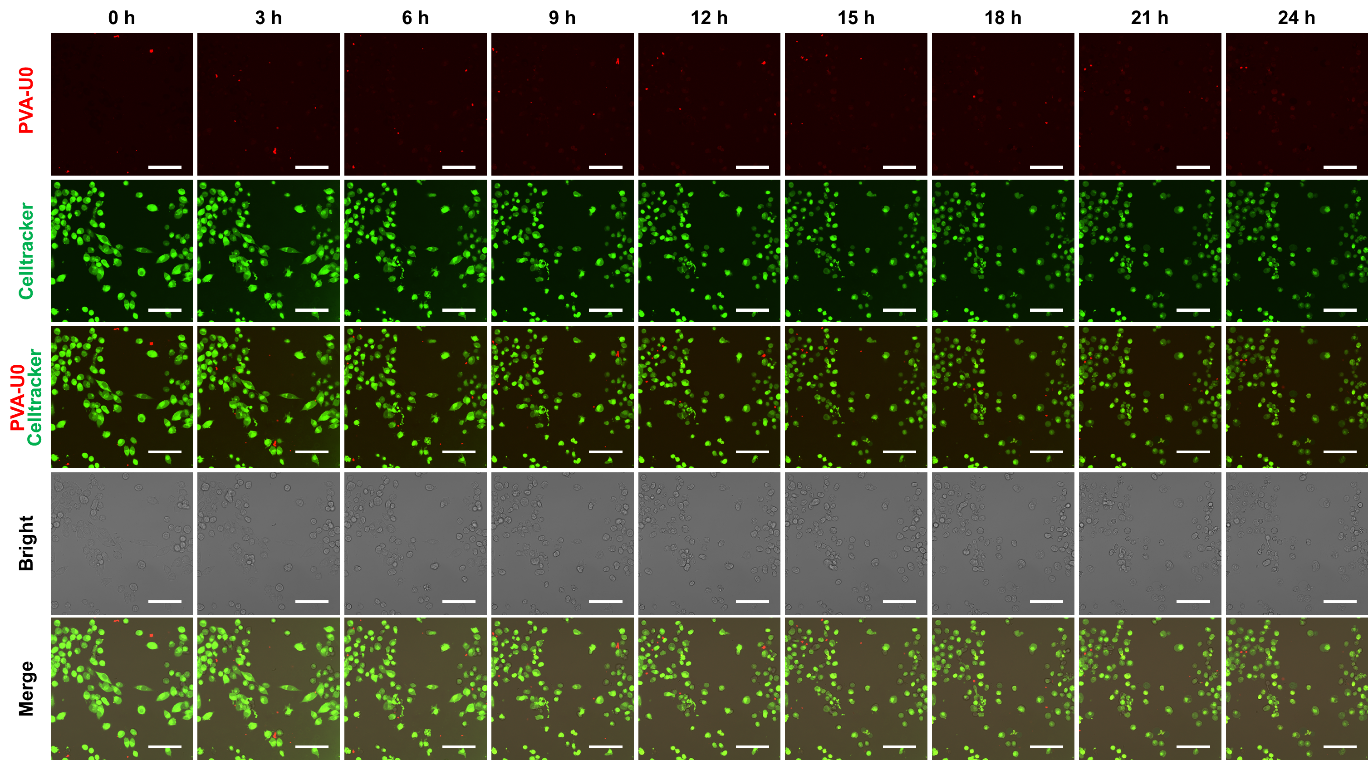

**Figure S20.** Confocal and bright filed images of MiaPaCa-2 cells treated with Celltracker and PVA-U0 at pH 6.5. Scale bar is 50 µm.

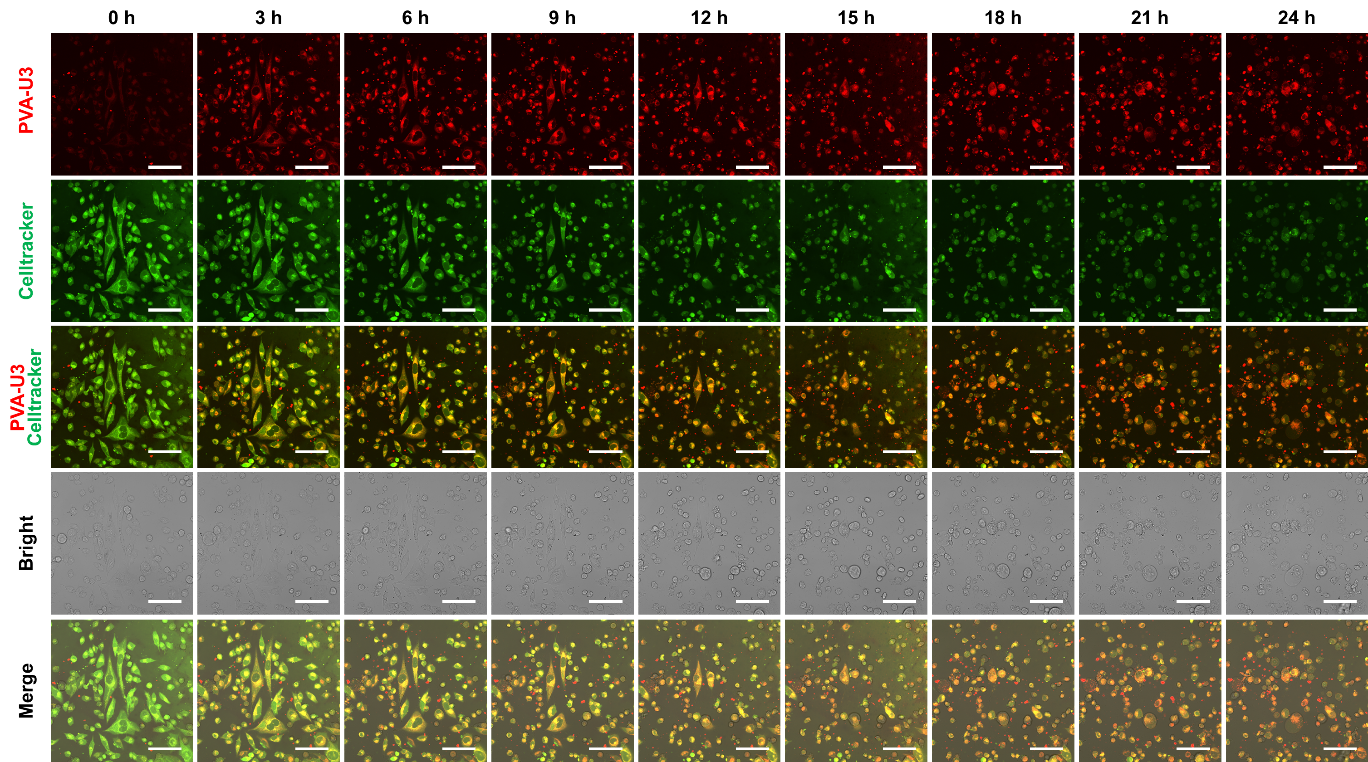

**Figure S21.** Confocal and bright filed images of MiaPaCa-2 cells treated with Celltracker and PVA-U3 at pH 7.4. Scale bar is 50 µm.

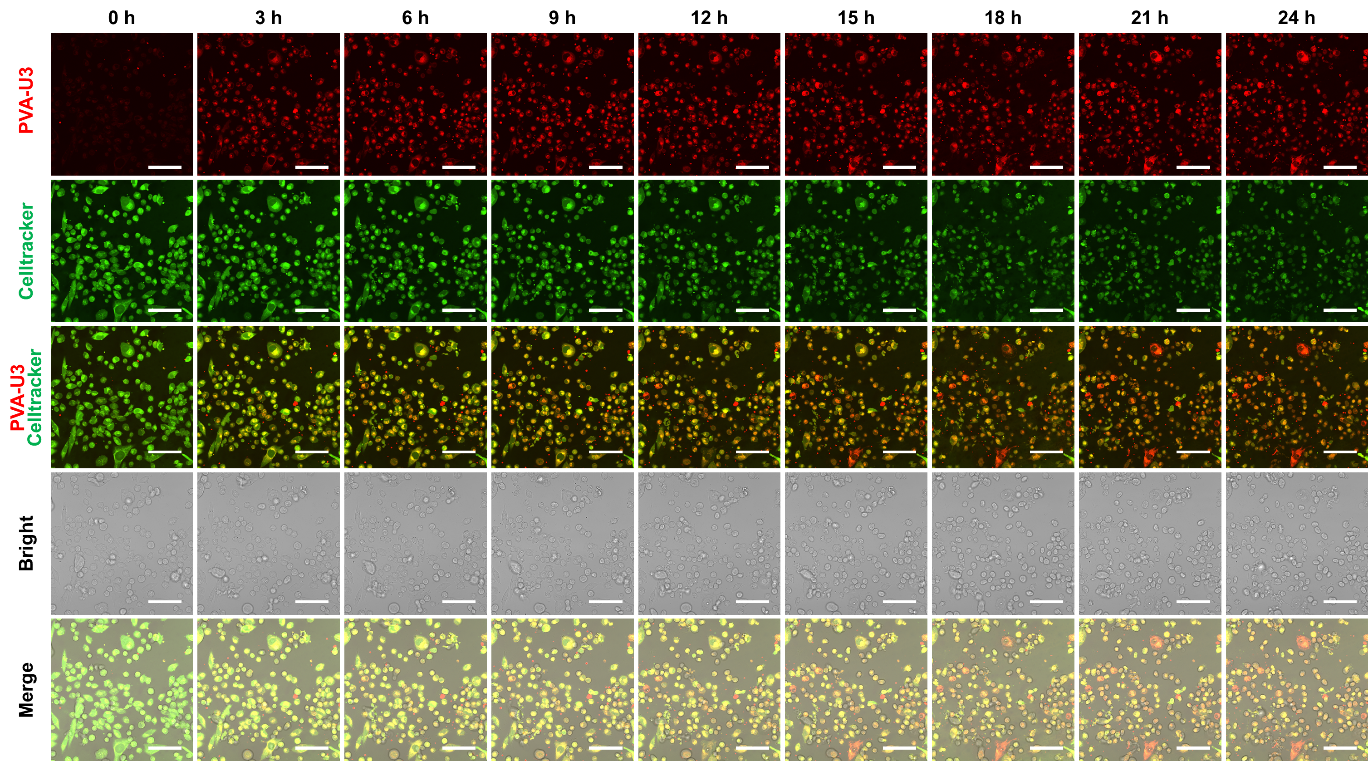

**Figure S22.** Confocal and bright filed images of MiaPaCa-2 cells treated with Celltracker and PVA-U3 at pH 6.5. Scale bar is 50 µm.

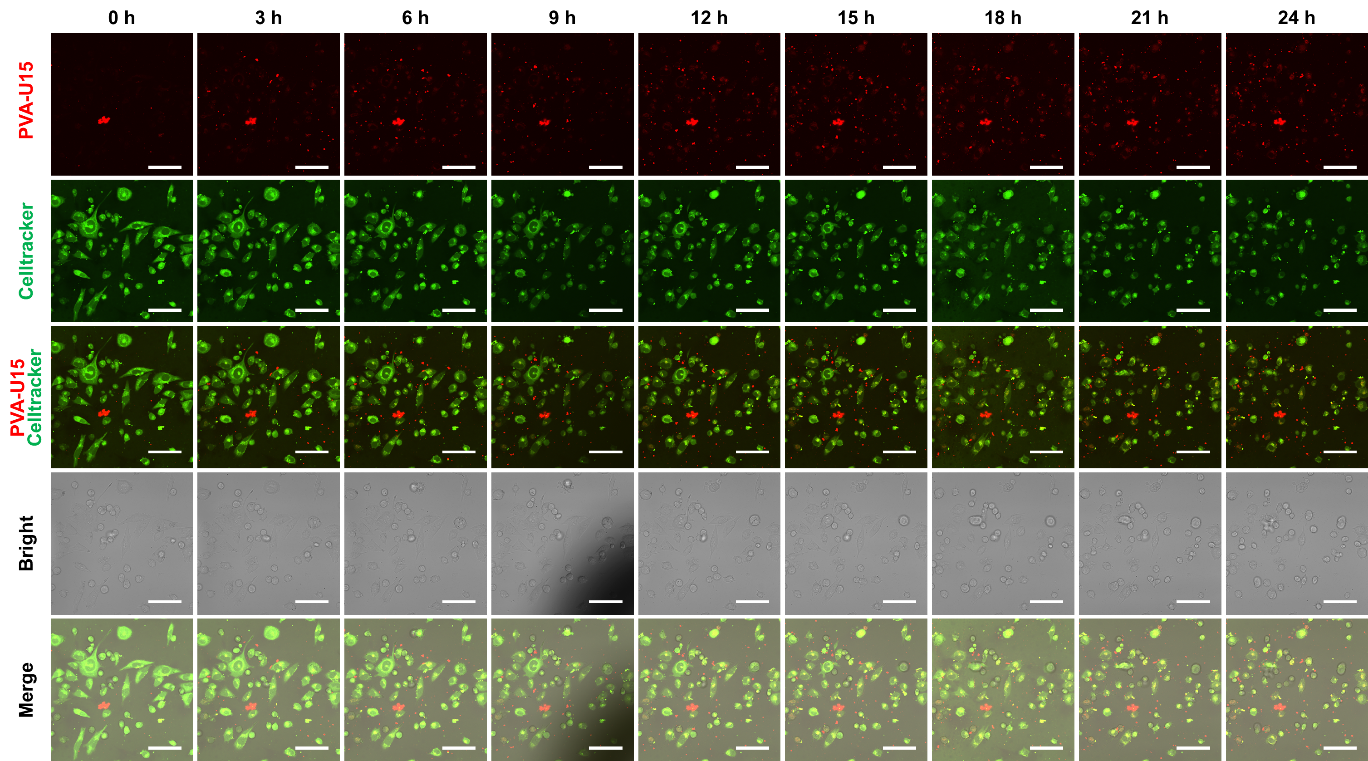

**Figure S23.** Confocal and bright filed images of MiaPaCa-2 cells treated with Celltracker and PVA-U15 at pH 7.4. Scale bar is 50 µm.

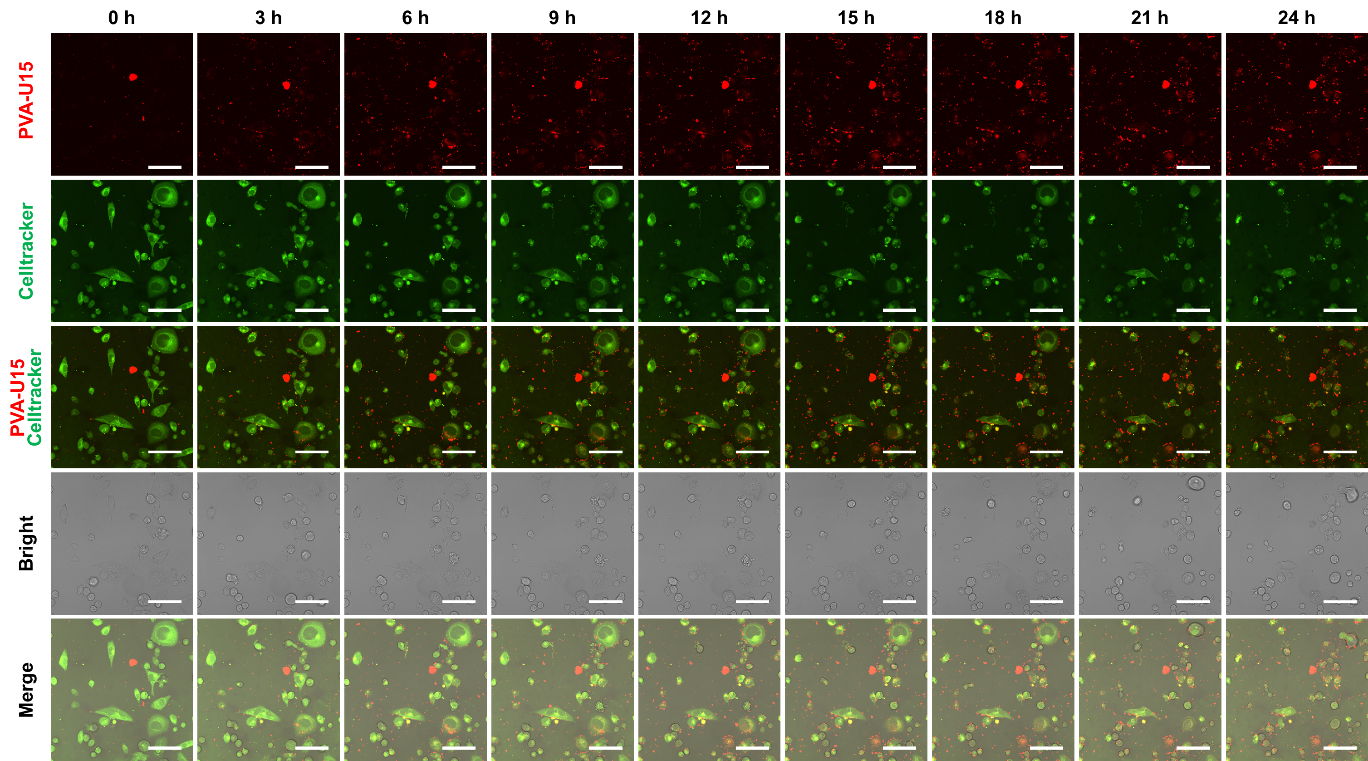

**Figure S24.** Confocal and bright filed images of MiaPaCa-2 cells treated with Celltracker and PVA-U15 at pH 6.5. Scale bar is 50 µm.

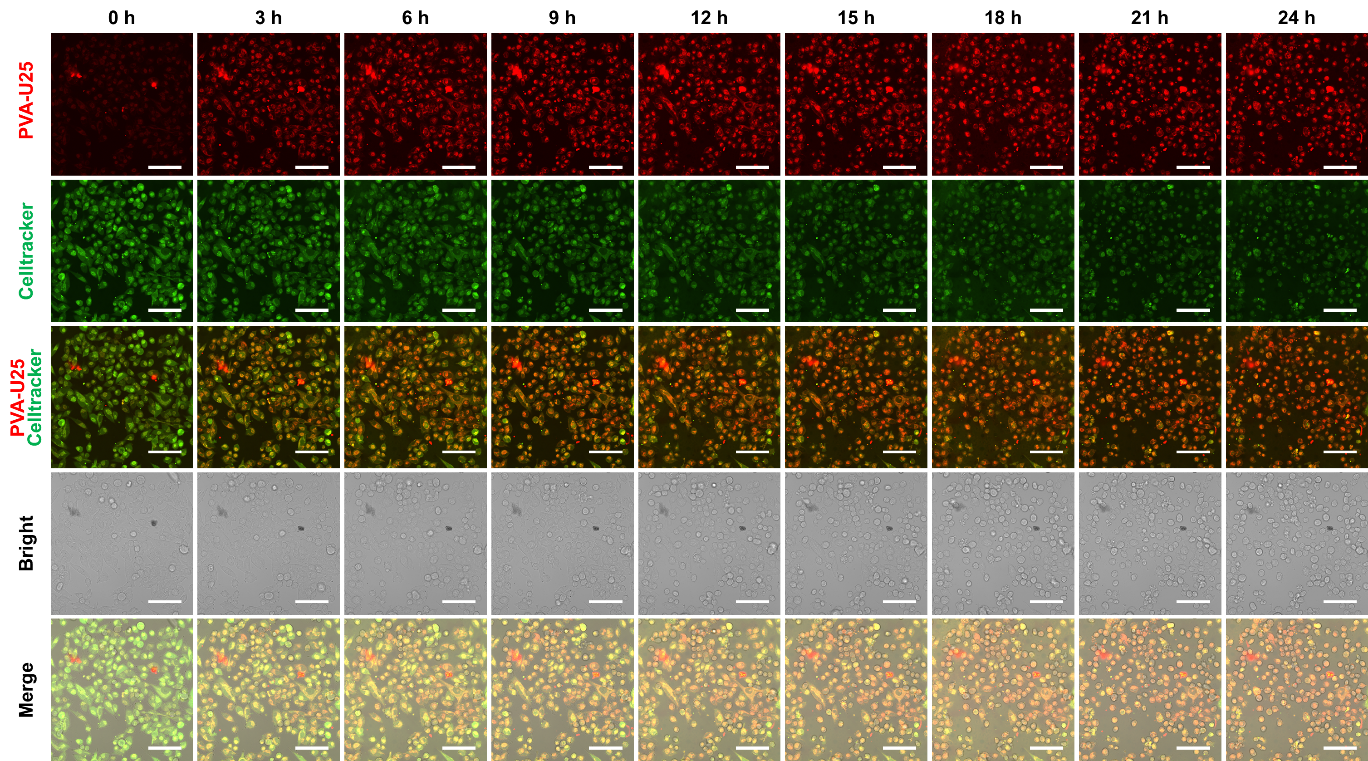

**Figure S25.** Confocal and bright filed images of MiaPaCa-2 cells treated with Celltracker and PVA-U25 at pH 7.4. Scale bar is 50 µm.

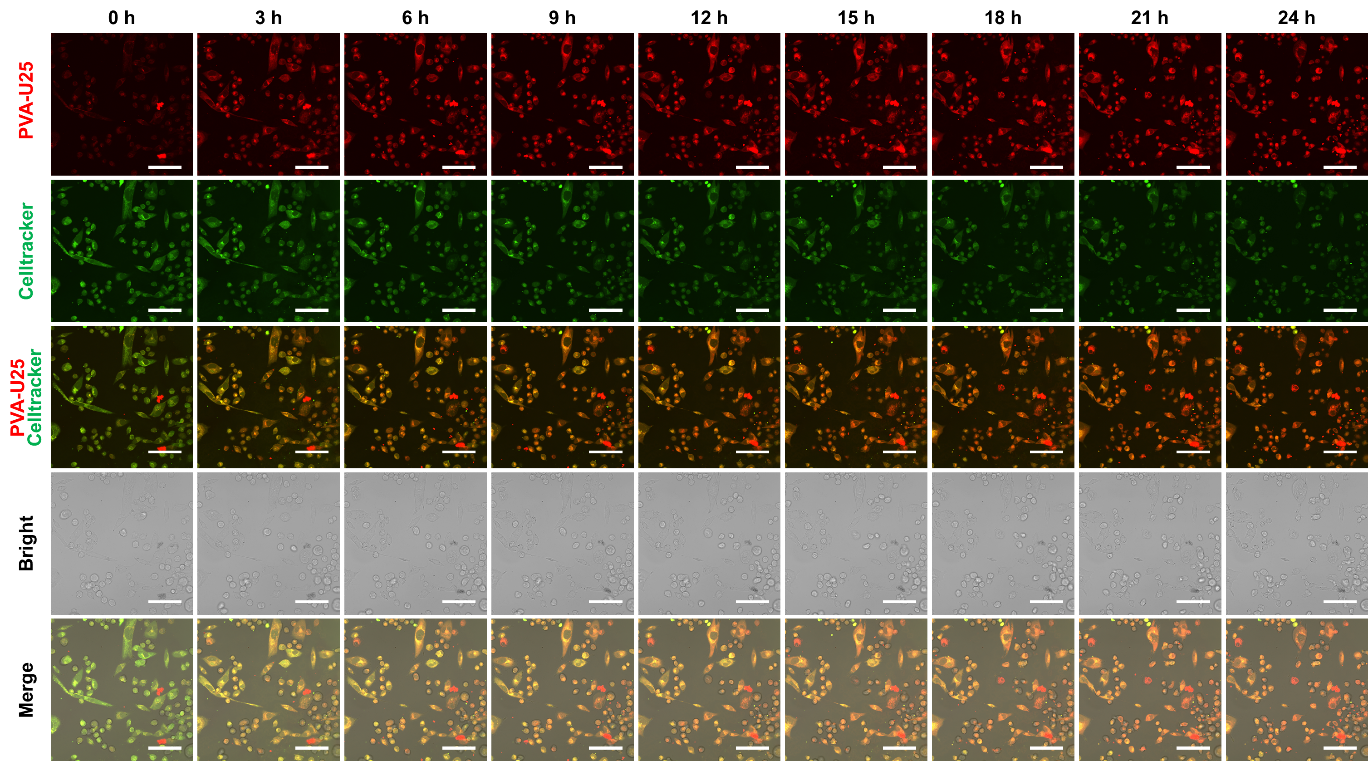

**Figure S26.** Confocal and bright filed images of MiaPaCa-2 cells treated with Celltracker and PVA-U25 at pH 6.5. Scale bar is 50 µm.

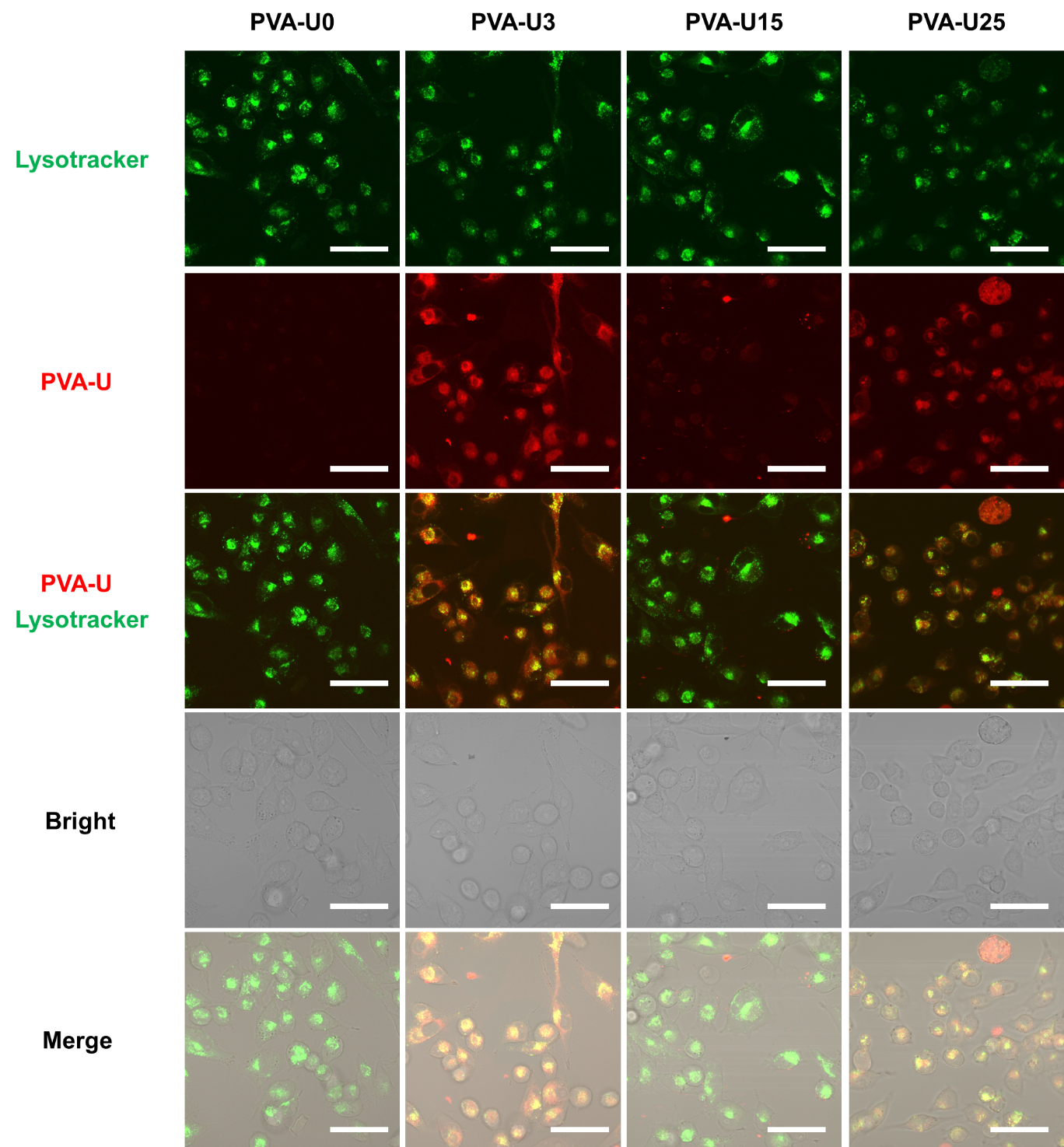

**Figure S27.** Confocal and bright filed images of MiaPaCa-2 cells treated with lysotracker and PVA-U at pH 7.4. Scale bar is 50 µm.

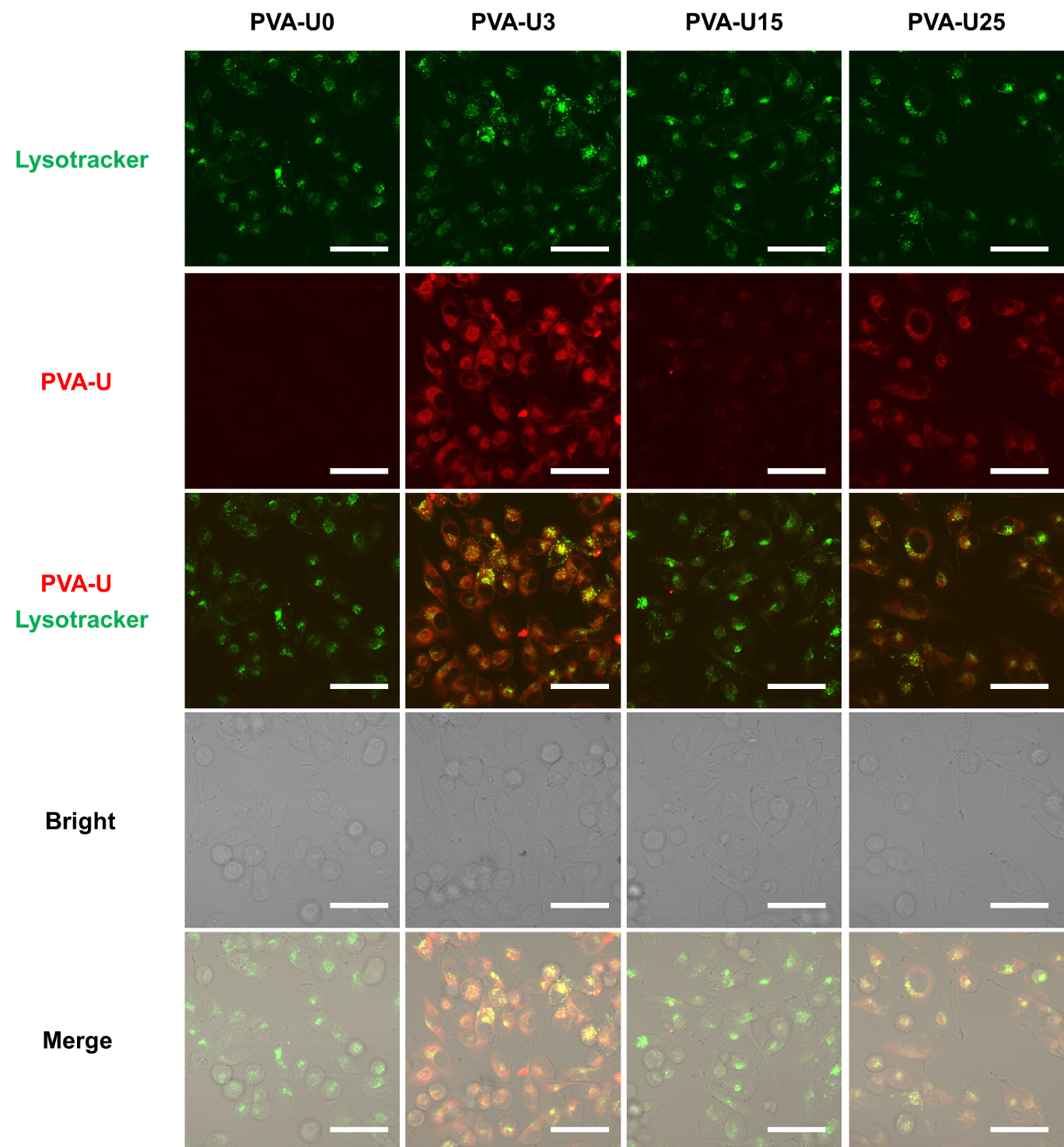

**Figure S28.** Confocal and bright filed images of MiaPaCa-2 cells treated with lysotracker and PVA-U at pH 6.5. Scale bar is 50 µm.

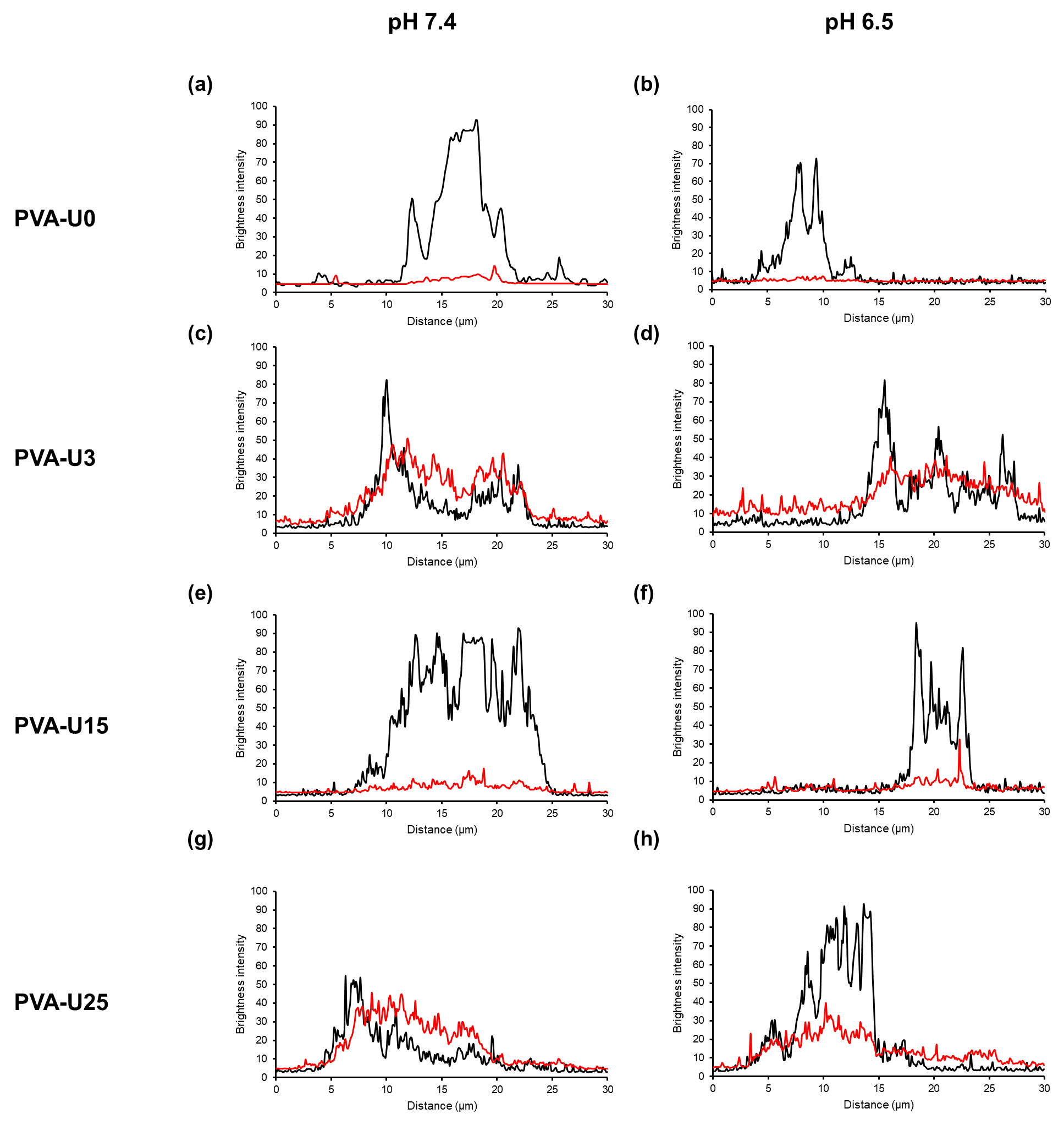
**Figure S29.** Line scan profiles of lysotracker (black) and PVA-U0 (a, b), PVA-U3 (c, d), PVA-U15 (e, f), and PVA-U25 (g, h) (red) at pH 7.4 and 6.5.

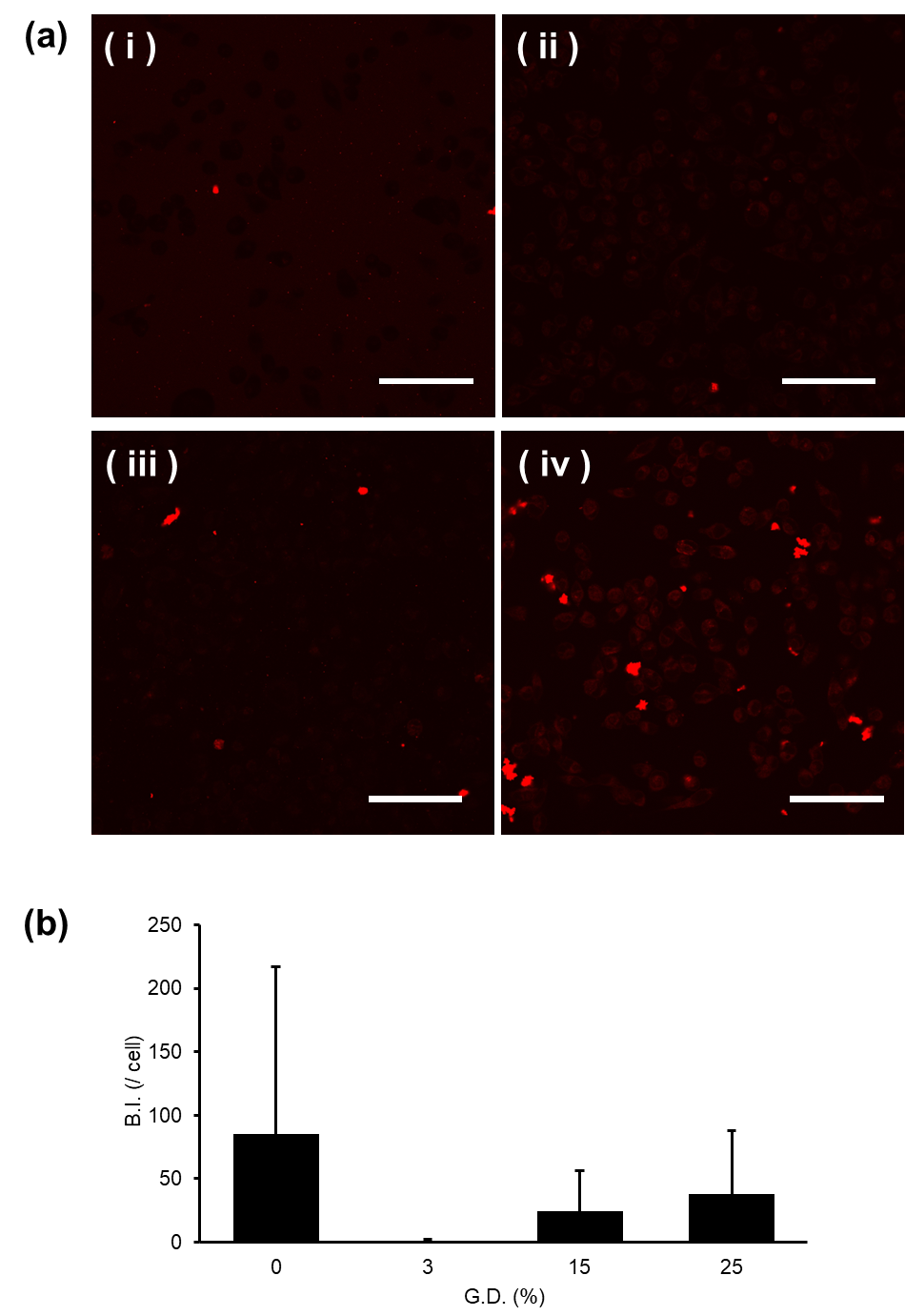

**Figure S30.** (a) Confocal images of MiaPaCa-2 cells treated with PVA-U0 (i), PVA-U3 (ii), PVA-U15 (iii), and PVA-U25 (iv) incubated for 4 h at 4 ^o^C. (b) Intracellular B.I. of PVA-U0, PVA-U3, PVA-U15, and PVA-U25 at pH 6.5.

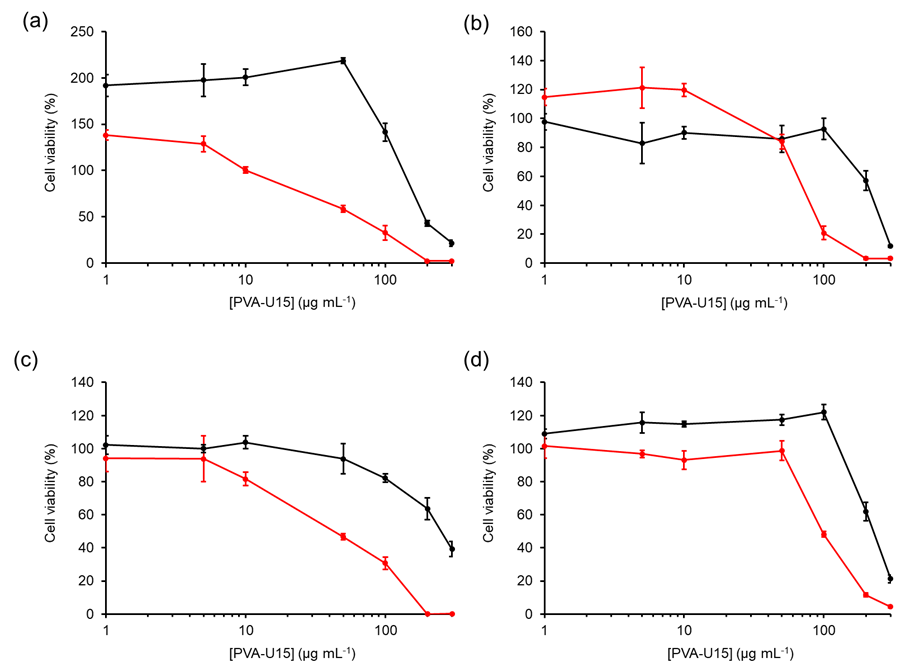

**Figure S31.** Cell viability of colon cancer (HT-29) cells (a), breast cancer (MDA-MB-231) cells (b), lung cancer (A549) cells (c), and normal human dermal fibroblast (NHDF) (d) treated with 10 µg mL^-1^ PVA-U15 at pH 7.4 (black) and pH 6.5 (red) after 24 h incubation
